# The Imposition of Value on Odor: Transient and Persistent Representations of Odor Value in Prefrontal Cortex

**DOI:** 10.1101/753426

**Authors:** Peter Y. Wang, Cristian Boboila, Philip Shamash, Zheng Wu, Nicole P Stein, L.F. Abbott, Richard Axel

## Abstract

The representation of odor in olfactory cortex (piriform) is distributive and unstructured and can only be afforded behavioral significance upon learning. We performed 2-photon imaging to examine the representation of odors in piriform and in two downstream stations, the orbitofrontal cortex (OFC) and medial prefrontal cortex (mPFC), as mice learned olfactory associations. In piriform we observed minor changes in neural activity unrelated to learning. In OFC, 30% of the neurons acquired robust responses to conditioned stimuli (CS+) after learning, and these responses were gated by context and internal state. The representation in OFC, however, diminished after learning and persistent representations of CS+ and CS− odors emerged in mPFC. Optogenetic silencing indicates that these two brain structures function sequentially to consolidate the learning of appetitive associations. These data demonstrate the transformation of a representation of odor identity in piriform into transient and persistent representations of value in the prefrontal cortex.

## INTRODUCTION

Most organisms have evolved a mechanism to recognize olfactory information in the environment and transmit this information to the brain where it must be processed to create an internal representation of the external world. This representation must translate stimulus features into appropriate behavioral responses. The olfactory sensory system does not merely represent the external world. Rather it interprets features of the world and combines them in higher cortical centers to construct representations that encode both the identity and value of different odors. Only two synapses intervene between the nose and the olfactory cortex, the piriform. Piriform cortex projects directly to higher order brain structures such as the orbitofrontal cortex (OFC) and medial prefrontal cortex (mPFC) (Chen et al., 2014; Diodato et al., 2016; Price, 1985). This shallow and well-characterized sensory pathway affords us the ability to identify the circuits that transform the identity of a sensory stimulus into representations of value that guide behavior.

Olfactory perception is initiated by the recognition of odorants by a large repertoire of receptors in the sensory epithelium (Buck and Axel, 1991; Godfrey et al., 2004; Zhang and Firestein, 2002). Individual sensory neurons in mice express only one of 1100 different receptor genes, and neurons that express the same receptor project with precision to two spatially invariant glomeruli in the olfactory bulb (Mombaerts et al., 1996; Ressler et al., 1993, 1994; Vassar et al., 1994). Thus, a transformation in the representation of olfactory information is apparent in the bulb where a dispersed population of active neurons in the sense organ is consolidated into a discrete spatial map of glomerular activity (Bozza et al., 2004). Each odorant activates a unique ensemble of glomeruli and the recognition of an odor requires integration of information from multiple glomeruli in higher olfactory centers.

The projection neurons of the olfactory bulb, the mitral and tufted cells, extend an apical dendrite into a single glomerulus and send axons to several telencephalic areas including significant input to piriform cortex (Price and Powell, 1970). Anatomic tracing reveals that axonal projections from individual glomeruli discard the spatial patterning of the bulb and diffusely innervate the piriform (Ghosh et al., 2011; Sosulski et al., 2011). Electrophysiologic and optical recordings demonstrate that individual odorants activate subpopulations of neurons distributed across the piriform without apparent spatial preference (Illig and Haberly, 2003; Iurilli and Datta, 2017; Poo and Isaacson, 2009; Rennaker et al., 2007; Stettler and Axel, 2009; Sugai et al., 2005; Zhan and Luo, 2010). Moreover, exogenous activation of an arbitrarily chosen ensemble of piriform neurons can elicit behaviors of contrasting valence dependent on learning (Choi et al., 2011). These observations are consistent with a model in which individual piriform cells receive convergent input from a random collection of glomeruli (Davison and Ehlers, 2011; Miyamichi et al., 2011; Stettler and Axel, 2009). In this model odor representations in piriform can only be afforded behavioral significance upon learning.

The piriform cortex sends projections to numerous brain regions including the amygdala, hippocampus, and prefrontal cortex, and is anatomically poised to accommodate the transformation of sensory representations into representations of value that can lead to appropriate behavioral output (Chen et al., 2014; Diodato et al., 2016; Johnson et al., 2000; Price, 1985; Schwabe et al., 2004). Neurons in orbitofrontal cortex (OFC) in both rodents and primates represent value but also encode other task variables including stimulus identity, motor action, confidence, internal state and task context (Feierstein et al., 2006; Gottfried et al., 2003; Kepecs et al., 2008; Lipton et al., 1999; Namboodiri et al., 2019; Padoa-Schioppa and Assad, 2006; Ramus and Eichenbaum, 2000; Schoenbaum and Eichenbaum, 1995; Schoenbaum et al., 1998, 1999; Thorpe et al., 1983; Tremblay and Schultz, 1999). Lesion experiments implicate OFC in updating learned information but these studies failed to reveal a role for OFC in simple associative learning (Bissonette et al., 2008; Burke et al., 2008; Chudasama and Robbins, 2003; Gallagher et al., 1999; Izquierdo et al., 2004; Ostlund and Balleine, 2007; Schoenbaum et al., 2002; Stalnaker et al., 2007). Medial prefrontal cortex (mPFC) has been implicated in simple associative learning and the remodeling of learned information (Birrell and Brown, 2000; Bissonette et al., 2008; Chudasama and Robbins, 2003; Ferenczi et al., 2016; Kim et al., 2017; Kitamura et al., 2017; Ostlund and Balleine, 2005; Otis et al., 2017). Recently, a neural representation of rewarded auditory stimuli was identified in both OFC and mPFC, and silencing of these brain structures elicited deficits in the acquisition and expression of learned behavior (Namboodiri et al., 2019; Otis et al., 2017).

We have performed two photon endoscopic imaging in piriform, OFC, and mPFC during appetitive associative conditioning to identify brain structures that exhibit changes in their neural representations upon olfactory learning (Barretto et al., 2009; Denk et al., 1990; Jung et al., 2004). Optogenetic silencing was then used to discern possible roles for these representations in associative conditioning. Imaging of neural activity in the piriform revealed that odor responses were sparse, selective, and unchanged by learning. Imaging of neural activity in the orbitofrontal cortex (OFC) revealed that 30% of OFC neurons acquired robust responses to conditioned (CS+) but not to unconditioned (CS−) odors during training. Moreover, these responses were gated by context and internal state. This representation in OFC diminished after learning, and persistent and non-overlapping representations of CS+ and CS− odors emerged in mPFC. Optogenetic silencing revealed that the OFC and mPFC appear to function sequentially in the learning of appetitive associations. These data demonstrate the transformation of a representation of odor identity in piriform into a transient representation of positive value in the OFC and then a persistent representation of positive and negative value in the mPFC.

## RESULTS

### Representation of Odor Identity in Piriform Cortex

We examined odor representations in piriform cortex while mice learned an appetitive odor discrimination task. Head fixed mice were exposed to two (CS+) odors that predicted a water reward delivered after a short delay and to two unrewarded (CS−) odors (Figure 1A). In separate trials the mice received a water reward (US) without prior odor delivery. After three to four training sessions, nearly all mice displayed anticipatory licking in response to the CS+ odors in over 90% of the trials (17 of 19 mice) and licked in fewer than 15% of the CS− trials (18 of 19 mice) (Figure 1B, 1C). We imaged neural activity by 2-photon microscopy during training in 6 mice expressing GCaMP6s in excitatory neurons in the piriform (Barretto et al., 2009; Chen et al., 2013; Denk et al., 1990; Jung et al., 2004; Madisen et al., 2015; Vong et al., 2011). We recorded the activity of 359 piriform neurons in six mice during one week of learning. Before learning, the four odors each activated an average of 16% of the piriform neurons (Figure 1F, S1B). Less than 4% of the neurons in piriform responded to water without prior odor exposure (Figure 1E, S1B). The neural responses after learning were largely unchanged (Figure 1D, 1F, S1A, S1B, S1F-H). The ensemble evoked by a given odor prior to training was significantly correlated with the ensemble evoked by the same odor after four days of task learning (Figure 1G). In a separate series of experiments, across-day correlations were similarly high after four days of passive odor exposure (Figure 1G). We also measured the correlation between the ensemble activities evoked by pairs of distinct odors. We found that the correlations between pairs of odor ensembles were low prior to learning and decreased even further after learning (Figure 1H, 1I; before learning: 0.50, after learning: 0.32, p < 0.001, Wilcoxon signed-rank test.). These results suggest that odor representations are stable but become slightly more discriminable across multiple training days.

**Figure 1.**
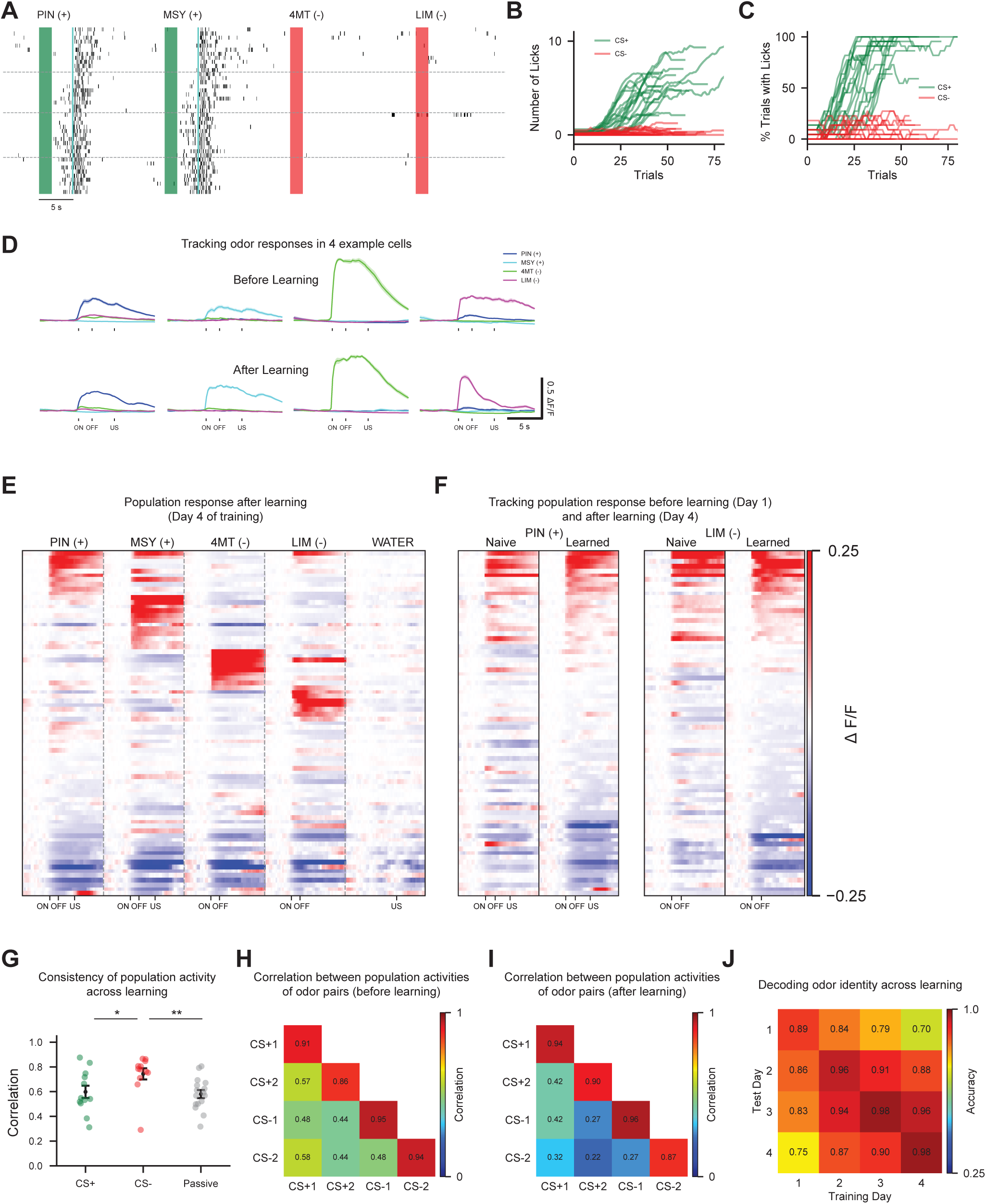
Odor representations in piriform cortex are stable during learning. (A) Anticipatory licking behavior in response to CS+ (green) but not CS− (red) odors after learning. Odor was presented for two seconds (green/red bars), followed by a three second delay before water delivery. Black rasters denote single lick events. Horizontal lines denote the 4 training sessions. Odors: PIN: pinene (CS+), MSY: methyl salicylate (CS+), 4MT: 4-methylthiazole (CS−), LIM: limonene (CS−). (B and C) Summary of training data for the appetitive odor discrimination task (n = 19 mice, combined across multiple imaging and behavioral experiments). Number of anticipatory licks (B) and percentage of trials with anticipatory licking (C) to CS+ and CS− odors. Green lines represent an average of the two CS+ odor trials for a single mouse and red lines represent averages of the two CS− odor trials for a single mouse. (D) Trial-averaged responses of 4 example piriform neurons to odors before learning (top, day 1 of training) and after learning (bottom, day 4 of training). ON: odor onset. OFF: odor offset. US: water delivery. Shading indicates ±1 SEM. (E) PSTH of piriform responses for one mouse. Cells are sorted by response amplitude to each of the 4 odors. Each row denotes a single cell’s trial-averaged responses to the four odors and water. For all PSTH plots, mean activity during the baseline period (5 seconds prior to odor delivery) is subtracted from each cell. See STAR Methods. (F) PSTH before learning and after learning for PIN (CS+) and LIM (CS−) from the same mouse as in (E). For each odor, responses are aligned across days and sorted by evoked amplitude after learning. (G) Pearson correlation was calculated between vectors encoding odor-evoked population activity before learning (day 1 of training) vs. after learning (day 4 of training), and day 1 vs. day 4 of passive odor exposure. Each dot represents across-day correlations for a single odor in one mouse. Correlation values were: CS+: 0.60, CS-: 0.74, passive: 0.58. N = 6 mice for odor learning, N = 4 mice for passive odor exposure. Green: CS+ odors. Red: CS− odors. Gray: passive. * P < 0.05, ** P < 0.01, Dunn’s test for multiple comparisons. Error bars indicate mean ±1 SEM. See STAR Methods. (H and I) Correlation of activity evoked by all odor pairs before (H) and after (I) learning. Correlations are calculated using the population activities for all pairs of odors in a given day and are averaged across the 6 imaged mice. Average correlation for all pairs of distinct odors before learning: 0.50, after learning: 0.32, p < 0.001, Wilcoxon signed-rank test. Diagonal entries represent self-correlations calculated after splitting shuffled trials of a given odor into two equal halves. See STAR Methods. (J) Accuracy of decoding the identities of the four tested odors from population activity within and across training days. Chance accuracy is 25%. Before learning (train and test on day 1): 0.89, after learning (train and test on day 4): 0.98, p = 0.04, Wilcoxon signed-rank test. 40 randomly chosen neurons per animal were used and values shown were averaged across 100 repetitions and 6 imaged mice. See STAR Methods. See also Figure S1.

These conclusions are further supported by decoding analysis (Figure S1C). A linear decoder trained on population activity prior to learning distinguished the identities of the four odors using population activity after learning (day 4 of training) with greater than 75% accuracy (Figure 1J). The odor ensembles became slightly more separable as training proceeded (Figure 1J), but these changes were qualitatively similar for both CS+ and CS− odor ensembles (Figure S1D, S1E), and also occurred upon passive odor exposure (Figure S1F-K). Thus, minor changes were observed in the representation of odors after training, but these changes were observed for both CS+ and CS− odors and were not dependent upon learning. These experiments suggest that changes in the representation of odor in the piriform do not reflect learning, and neural instantiations of learning must occur downstream.

### A Representation of Value in Orbitofrontal Cortex

The piriform cortex sends axons to numerous brain regions, with an extensive projection to orbitofrontal cortex (Chen et al., 2014; Price, 1985). We therefore asked whether appetitive odor learning elicits changes in the representation of odors in OFC. We imaged the activity of 364 OFC neurons in 5 animals across multiple training days. Before learning, the four odors each activated an average of 12% of the neurons in OFC (Figure 2A, S2B). The responses were non-selective, inconsistent, and low in amplitude (Figure 2A, 2C, S2A). 16% of imaged neurons responded to water, the unconditioned stimulus (Figure S2B). In contrast to piriform cortex, we could not discern a representation of odor identity in OFC prior to learning (see below).

**Figure 2.**
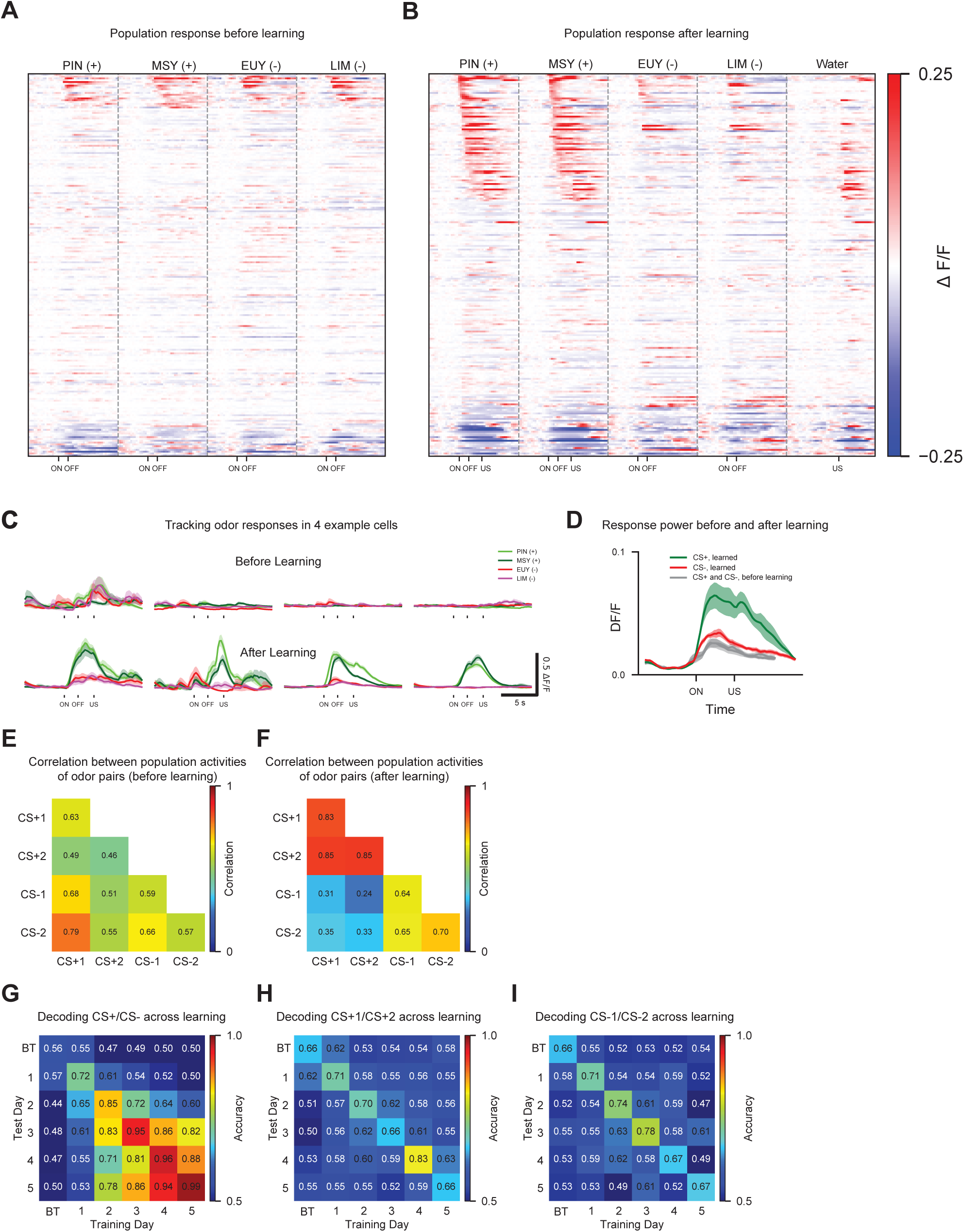
A CS+ representation emerges in the OFC after learning. (A and B) PSTH of OFC responses for all imaged mice (n=5) before (A) and after (B) learning. Odors: PIN: pinene (CS+), MSY: methyl salicylate (CS+), EUY: eucalyptol (CS−), LIM: limonene (CS−). Responses before and after learning are not aligned but sorted for each panel by response amplitude to CS+ odors. Due to differences in experimental conditions, imaging data from 1 mouse could not be combined with the 4 other mice in (A), so 4 mice are shown in (A). (C) Trial-averaged responses of 4 example OFC cells to odors before learning (top) and after learning (bottom). ON: odor onset. OFF: odor offset. US: water delivery. Here and below, shading indicates ±1 SEM. (D) Average response power of OFC neurons to CS+ and CS− odors before learning (all odors: gray), and after learning (CS+: green, CS-: red). N=5 mice. See STAR Methods. (E and F). Within-day correlations between odor ensembles before learning (E) and after learning (F). Average correlation between the population activities evoked by CS+1 and CS+2 before learning: 0.49, after learning: 0.85, p = 0.04, Wilcoxon signed-rank test. Correlation between all CS+/CS− odor pairs before learning: 0.63, after learning: 0.30, p < 0.001, Wilcoxon signed-rank test. (G-I). Accuracy of decoding predictive value (CS+ odors vs. CS− odors, G), CS+ identity (CS+1 vs. CS+2, H), and CS− identity (CS-1 vs. CS-2, I) from OFC population activity within and across training days. 40 randomly chosen neurons per animal were used. Values shown are averaged across 100 repetitions and 5 imaged mice. Chance accuracy is 50% for all three conditions. BT: before training. See also Figure S2 and S3.

We observed a striking change in the neuronal response to CS+ odors as learning proceeded (Figure 2B, 2C). After learning, 30% of the OFC neurons acquired consistent, high amplitude responses to each of the two CS+ odors (Figure S2B). 75% of neurons responsive to one CS+ odor also responded to the second CS+ odor (Figure S2C). Moreover, the amplitude and duration of responses to the two CS+ odors in a given neuron were similar (Figure S2D, S2E). 64% of the CS+ responsive neurons were not activated in water-only trials, demonstrating that the majority of these CS+ responses did not result from the activation of a motor program (Figure 2B). After learning, CS− odors continued to elicit sparse, inconsistent, and low amplitude responses, similar to the responses observed prior to training (Figure 2B-D, S2B). These observations suggest projections from the CS+ representation in piriform to the OFC are reinforced during learning.

The mean excitatory response amplitude (which we call the response power) evoked by CS+ odors increased almost three-fold (286%) after learning, with only a minor (138%) change in response power to CS− odors (Figure 2D). Moreover, the population response to the two CS+ odors became highly correlated after learning, whereas the population response to the CS+ and CS− odors became less correlated (Figure 2E, 2F).

We performed decoding analysis to further examine the effect of learning on the OFC representation. A linear decoder trained on population responses prior to training decoded odor identity in the OFC at near chance levels (41% accuracy, chance is 25%). A decoder trained on population responses after learning distinguished between rewarded and unrewarded odors with greater than 95% accuracy (Figure 2G, S2F). In contrast, a decoder trained to distinguish between the identities of the two CS+ odors in OFC performed at close to chance level (Figure 2H, S2F). A decoder also failed to distinguish between the identities of the two CS− odors after learning (Figure 2I, S2F). This is in accord with our observation that the population activities between the two CS+ odors are highly correlated. These data suggest that the representation of odor identity encoded in piriform is discarded in the OFC and transformed into a representation of positive value by learning.

### The OFC Representation Reflects Changes in Value

If the value of an odor changes, the representation of value in OFC should also change (Roesch et al., 2007; Schoenbaum et al., 1999; Thorpe et al., 1983). We therefore recorded the neural responses in OFC during reversal learning. Mice were first trained with 2 CS+ and 2 CS− odors in the appetitive learning task, and the odor reward contingencies were then reversed. After reversal, the mice displayed anticipatory licking to the old CS− odors (CS+ upon reversal) and suppressed anticipatory licking to the old CS+ odors (CS− upon reversal) after 30 trials (Figure S4A). Prior to reversal, imaging revealed that 30% of the neurons were more responsive to CS+ than CS− odors (Figure 3A, 3B, see STAR Methods). After reversal learning, 91% of CS+ responsive neurons diminished their response to the old CS+ odors, and 68% of these neurons were now activated by the new CS+ odors (S4B, S4C). As a consequence, 28% of OFC neurons are now more responsive to the new CS+ than CS− odors (Figure 3B, S3B). We also analyzed the strength of the odor-evoked responses during reversal learning at the level of neuronal populations. The response power to the old CS+ odors diminished three-fold upon reversal (Figure 3C, 3D, green) whereas the response power to the old CS− odors increased three-fold upon reversal (Figure 3C, 3D, red). The observation that the same cells diminished their responses to the old CS+ odors and responded to the new CS+ odors after reversal (Figure 3A, S4B, S4C) indicates that these neurons encode value rather than odor identity.

**Figure 3.**
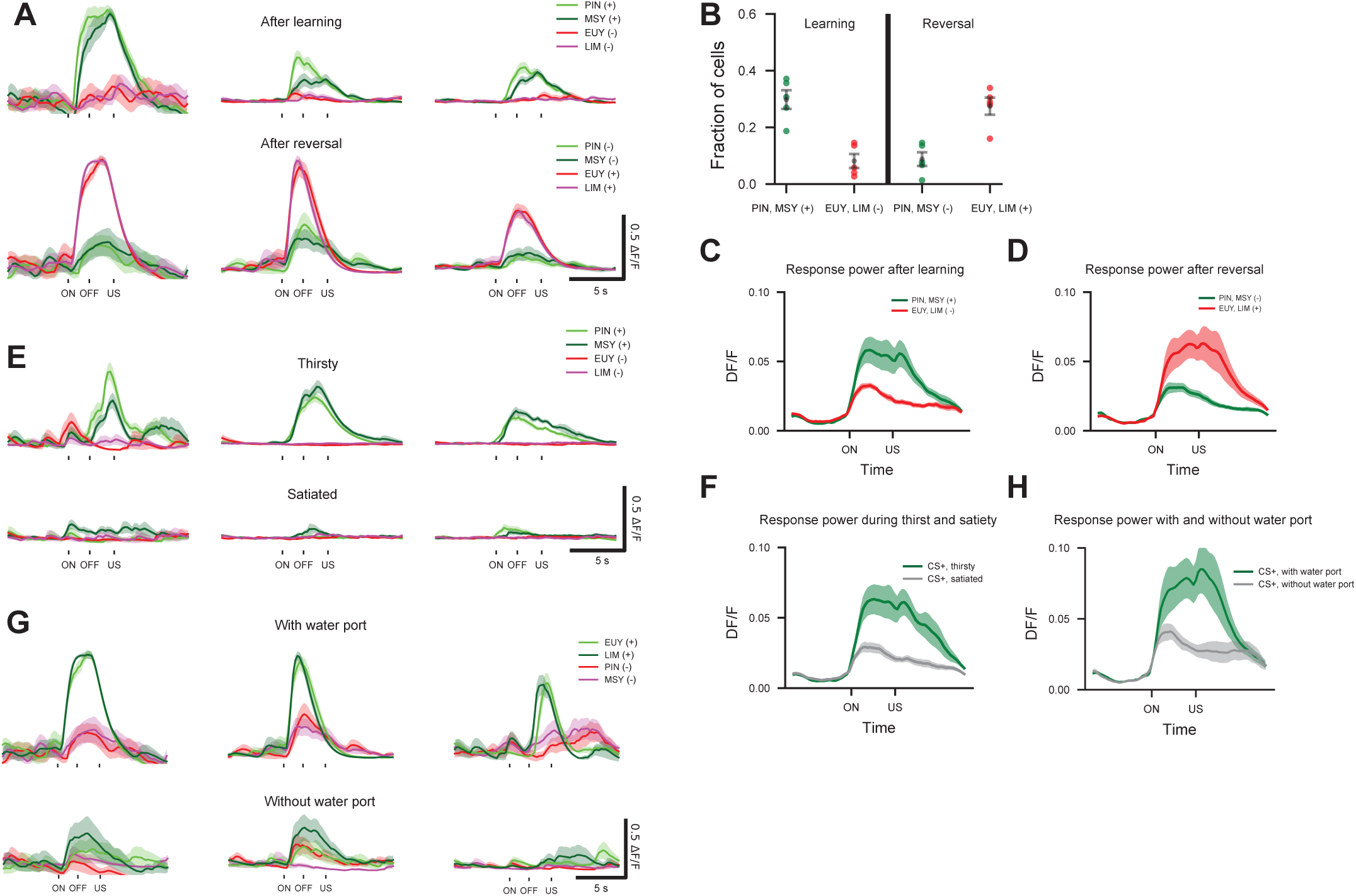
The CS+ representation in OFC is sensitive to internal state and external context. (A) Trial-averaged responses of 3 example OFC cells after learning (top) and after reversal (bottom). PIN and MSY (CS+) were rewarded during discrimination learning, but not during reversal. EUY and LIM (CS−) were not rewarded during discrimination learning, but were rewarded during reversal. Here and below, shading indicates ±1 SEM. (B) Fraction of neurons that are more responsive either to CS+ or to CS− odors after learning and after reversal for 5 mice. After learning CS+: 0.30, CS-: 0.08. After reversal CS+: 0.09, CS-: 0.27. Error bars indicate mean ±1 SEM and dots indicate individual animals. See STAR Methods. (C and D) Average response power of OFC neurons to CS+ (green) and CS− (red) odors after learning (C) and after reversal (D). (E) Trial-averaged responses of 3 example OFC cells in an animal that was thirsty (top) and then immediately satiated (bottom). (F) Average response power of OFC population to CS+ odors in thirsty (green) and satiated mice (gray). N = 5 mice. (G) Trial-averaged responses of 3 example cells when the water port was present (top) or absent (bottom). (H) Average response power of OFC population to CS+ odor when water port was present (green) or absent (gray). N=4 mice. See also Figure S3 and S4.

The value of a sensory stimulus should be contingent on internal state and context (Allen et al., 2019; Critchley and Rolls, 1996). CS+ odors predict water reward, an outcome of value to a thirsty mouse but of diminished value to a water-sated mouse. We therefore asked whether the representation of CS+ odors in OFC differs in thirsty and satiated mice. After appetitive learning, the mice were provided water. After satiation, the mice no longer displayed anticipatory licking to CS+ odors and rarely collect water when it is delivered (licking in less than 10% of trials) (Figure S4D). Imaging in the OFC revealed that prior to satiation, 30% of neurons responded to CS+, but 95% of these neurons were either no longer responsive or were significantly attenuated after satiation (Figure 3E, S3A). At a population level, the response power to the CS+ odors was more than 2.5-fold higher in thirsty mice (Figure 3F).

We also imaged mice for which the behavioral context was altered by removal of the water port. Under these conditions, water is not obtainable, and the value of the CS+ odor is presumably eliminated. Removal of the water port suppressed anticipatory licking to CS+ odors in less than three odor presentations (video recordings during imaging). Neuronal responses to the CS+ odors were either eliminated or significantly attenuated in 81% of the CS+ responsive neurons (Figure 3G, S3C). The response power to the CS+ odors was more than two-fold higher before water port removal (Figure 3H). Thus, changes in internal state and context that diminished the value of water reward correlated with a significant attenuation in the activity of the CS+ ensemble, providing further evidence that this OFC representation encodes value.

### The Role of the OFC Representation in Associative Learning

We next performed optogenetic silencing to ask whether the OFC contributes to the learning of an appetitive association. AAV encoding either halorhodopsin or the red-shifted halorhodopsin Jaws was injected bilaterally into OFC (Chuong et al., 2014; Gradinaru et al., 2008). Electrophysiological recording sessions using a 32-channel extracellular optrode array demonstrated that photostimulation results in over 4-fold inhibition in spontaneous activity in mice expressing Jaws and 8-fold inhibition in mice expressing halorhodopsin (Figure S5) (Royer et al., 2010). Silencing of OFC during training was initiated two seconds prior to odor delivery and extended for two seconds beyond the time of water delivery. Mice that experienced OFC inhibition exhibited significant learning deficits (Figure 4A-F). The 9 silenced mice either did not lick consistently to the CS+ odors or licked indiscriminately to CS+ and CS− odors, or both. The number of trials to criterion (anticipatory licking in over 80% of CS+ odor trials) was two-fold higher in OFC silenced mice than in control mice (Figure 4A, 4B). In addition, the number of trials required to suppress licking to CS− odors (anticipatory licking in less than 20% of CS− odor trials) was four-fold higher in OFC silenced mice than in control mice (Figure 4C, 4D). Moreover, 5 of 9 mice failed to reach criterion and were unable to discriminate between CS+ and CS− odors even after 100 presentations of each odor within 8-10 training sessions (Figure 4E). Both control and silenced mice exhibit robust licking upon water delivery, suggesting that mice with OFC inhibition were highly motivated to acquire water reward (Figure 4F). Thus, the neural representation of predictive value in OFC participates in the efficient acquisition of appetitive associations.

**Figure 4.**
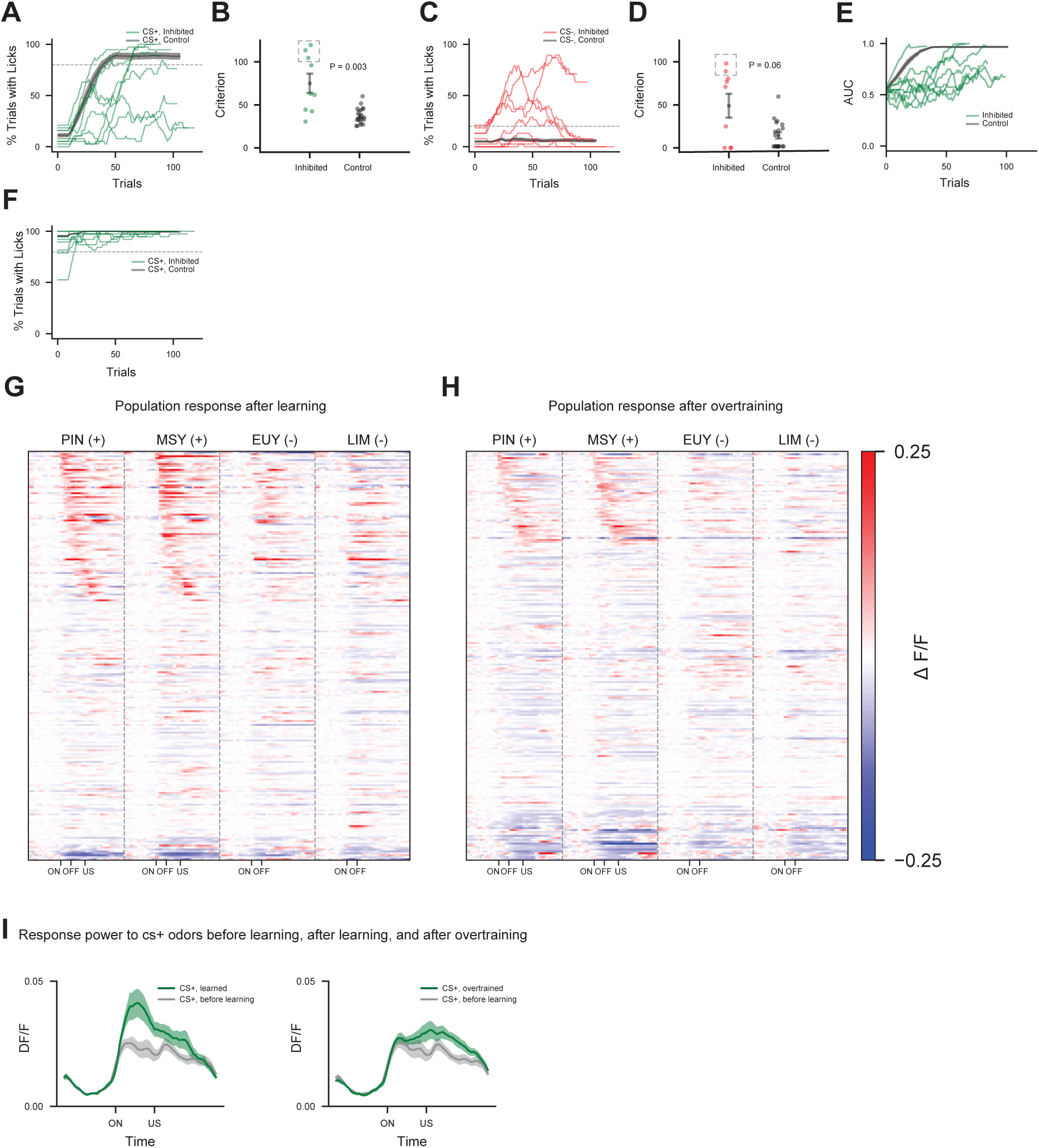
OFC is necessary for associative learning and the odor representation in OFC peaks during learning but diminishes after over-training. (A-F). Optogenetic silencing of OFC during appetitive learning. OFC inhibition (n=9 mice) was accomplished with either mice expressing Jaws (n=4) or halorhodopsin (n=5). Control animals (n=19) are pooled across conditions. See STAR Methods for inhibition protocol. (A) Fraction of trials with anticipatory licks to the CS+ odors. Dotted line indicates criterion for learning to lick to CS+ odors (>80% of trials with anticipatory licking). Here and below, shading indicates ±1 SEM for control animals. (B) Summary of trials to criterion for licking to CS+ odors in OFC silenced and control mice. Inhibited (green): 75 trials, control (gray): 37 trials, p = 0.003, ranksum test. Three inhibited mice did not reach criterion at the end of training (dotted square), and trials to criterion for these mice was defined as the last trial of training. Here and below, error bars indicate mean ±1 SEM and dots indicate individual animals. See STAR Methods. (C) Fraction of trials with anticipatory licks to the CS− odors. Dotted line indicates criterion for suppression of licking to CS− odors (<20% of trials with anticipatory licking). (D) Summary of trials to criterion for suppression of licking to CS− odors in OFC silenced and control mice. Inhibited (red): 49 trials, control (gray): 13 trials, p = 0.06, ranksum test. Two inhibited mice did not reach criterion at the end of training (dotted square), and trials to criterion for these mice was defined as the last trial of training. (E) Discriminability of licking to CS+ and CS− odors. AUC (area under ROC curve) was calculated by comparing the distribution of anticipatory licks in CS+ trials to that in CS− trials over a moving average window of 20 trials. An AUC of 0.5 indicates zero discriminability between licks to CS+ and CS− odors; an AUC of 1.0 indicates complete discriminability with more licking to CS+ than CS− odors. Green: inhibited, gray: controls. (F) Percentage of trials with collection licks to CS+ (green) odors. Green: inhibited mice. Gray: control mice. Collection licks are defined as the number of licks during the 1 second after water delivery. (G and H) PSTHs of OFC responses for all mice (n=3) to CS+ (PIN and MSY) and CS− (EUY and LIM) odors after learning (G) and after over-training (H). Responses after learning and after over-training are not aligned but sorted for each panel by response amplitude to CS+ odors. (I) Left: average response power of OFC neurons to CS+ odors before learning (grey) and after learning (green). Right: before learning (grey) and after over-training (green). See also Figure S5.

### The OFC Representation Diminishes After Learning

The CS+ representation in OFC was strongest after 3 to 4 days of training, a time when behavioral performance plateaus, but diminished at later times despite the persistence of learned behavior. We therefore performed imaging experiments in a new cohort of mice for longer periods extending up to 9 training sessions (Figure 4G, 4H). The response power of the CS+ representation was maximal at 3 to 4 days of training and declined to amplitudes observed prior to training after 6-9 days (Figure 4I). The observation that the CS+ representation in the OFC diminished whereas the behavior persisted suggests that OFC may participate in the acquisition of appetitive associations, but is no longer required after initial learning.

We considered the possibility that our olfactory association task may involve distinct phases of learning with only the initial phase dependent on OFC. We considered a behavioral model in which mice first learn that odor predicts water, and in a second phase of learning, acquire the ability to discriminate which odors predict reward. We therefore implemented a head-fixed associative learning task consisting of two phases, pre-training and discrimination (Figure 5A). This task is similar to learning paradigms in freely moving mice that require pre-training for task acquisition, but the role of specific brain regions in pre-training in these behavioral experiments has not been examined (Bissonette et al., 2008; Burke et al., 2008; Izquierdo et al., 2004; Schoenbaum et al., 1999, 2002, 2003; Stalnaker et al., 2007). In the pre-training phase of our new task a single odor was paired with water delivery. After mice successfully learned that odor predicts reward, a discrimination phase was initiated in which two new CS+ and two CS− odors were presented. This two-phase learning paradigm was conducted in cohorts of mice that express either Jaws or YFP in neurons in the OFC. OFC silencing in mice expressing Jaws impaired learning in the pretraining phase, with anticipatory licking requiring an average of 89 trials compared with 57 trials required by control mice (Figure 5B, 5C).

**Figure 5.**
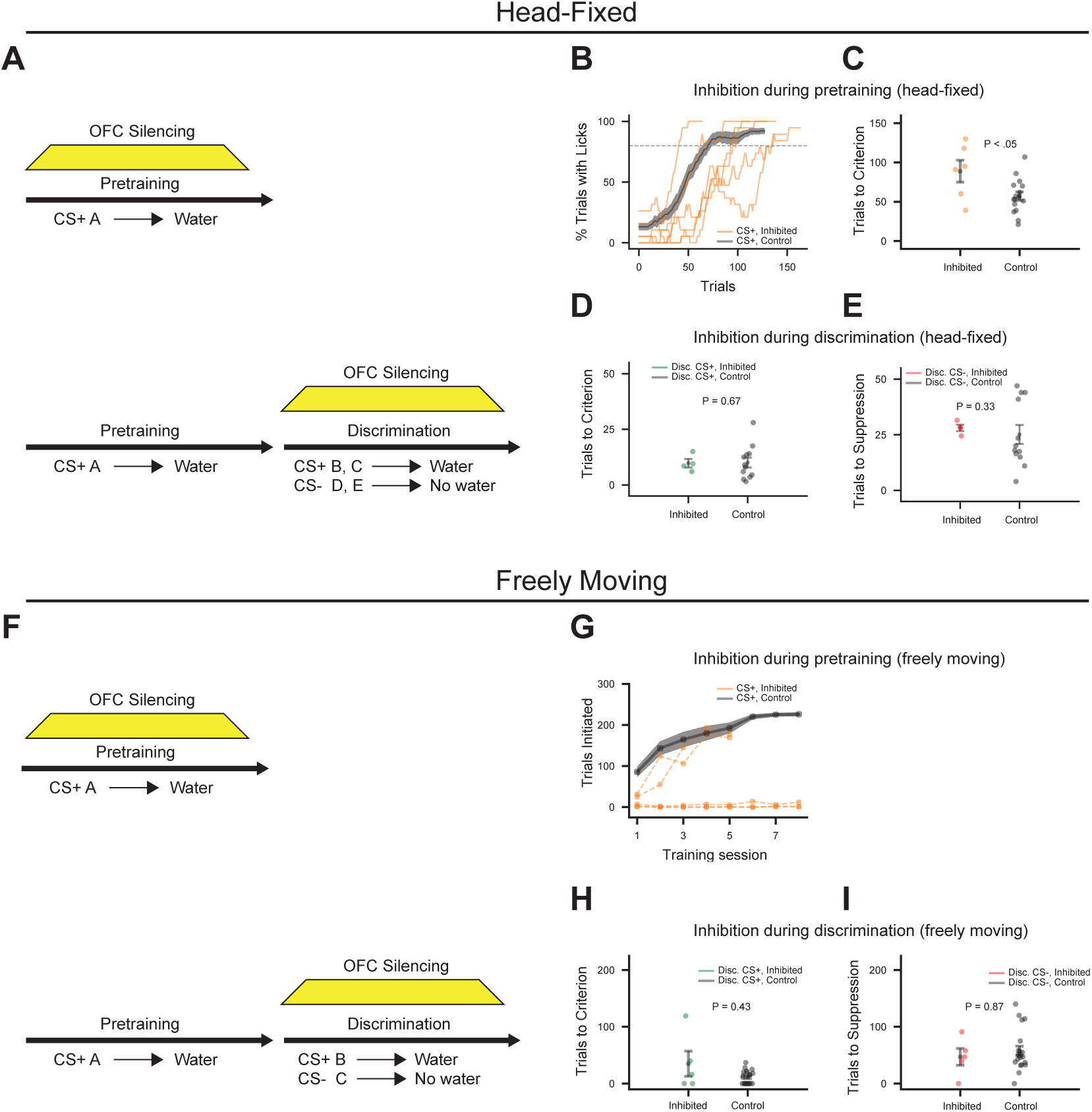
OFC is necessary for initial learning. (A) Schematic of optogenetic silencing of OFC in a head-fixed task. (B and C) OFC silencing during the pre-training phase of the two-phase task in head-fixed mice. Mice expressing Jaws: n=6, control mice: n=16. Control animals are pooled across conditions (see STAR Methods). (B) Percentage of trials with anticipatory licks to the pretraining CS+ odor (Mice expressing Jaws: orange, control: gray). Here and below, shading indicates ±1 SEM for control animals. (C) Trials to criterion for licking to the pretraining CS+ odor. Mice expressing Jaws (orange): 89 trials, control (gray): 57 trials, p = 0.04, ranksum test. Here and below, error bars indicate mean ±1 SEM and dots indicate individual animals. (D and E) OFC silencing during the discrimination phase of the two-phase task in head-fixed mice. Mice expressing Jaws: n=4, control mice: n= 12. (D) Trials to criterion for licking to CS+ odors. Mice expressing Jaws (green): 10 trials, controls (gray): 10 trials, p=0.67, ranksum test. (E) Trials to criterion for suppression of licking to CS− odors. Mice expressing Jaws (red): 25 trials, controls (gray): 28 trials, p = 0.33, ranksum test. We noted that mice expressing Jaws rapidly learned to suppress licking to CS− odors after 15 trials on the first day of discrimination training but may have a deficit in memory. (F) Schematic of optogenetic silencing in a freely moving task. (G) OFC silencing during the pre-training phase of the two-phase task in freely moving mice. Mice expressing halorhodopsin: n=5, orange; mice expressing YFP: n=21, gray. YFP animals are pooled across conditions. Number of trials initiated during pre-training is plotted as a function of training days. 3 of 5 mice with OFC inhibition failed to initiate trials. (H and I) Same as D, E for freely moving animals. Mice expressing halorhodopsin: n=5, mice expressing YFP: n= 21. (H) Trials to criterion for licking to CS+ odors. Mice expressing halorhodopsin (green): 35 trials, mice expressing YFP (gray): 11 trials, p=0.43, ranksum test. (I) Trials to criterion for suppression of licking to CS− odors. Mice expressing halorhodopsin (red): 47 trials, mice expressing YFP (gray): 58 trials, p = 0.87, ranksum test. See also Figure S6.

We next examined the role of OFC in the discrimination phase of the two-phase odor learning task. Mice expressing either Jaws or YFP in the OFC were pretrained in the absence of inhibition. After mice have successfully learned that odor predicts reward, anticipatory licking was observed in response to both the CS+ and CS− odors at the start of training (Figure S6A-D). This suggests that, during pretraining with a single CS+ odor, mice learn to generalize, associating all odors with reward. Control mice enhanced licking to the CS+ odors after 10 trials and suppressed licking to the CS− odors after 25 trials (Figure 5D, 5E, gray). Photoillumination of the OFC during the discrimination phase in mice expressing Jaws did not impair discrimination learning. Licking to CS+ odors (Figure 5D) and suppression of licking to CS− odors (Figure 5E) were similar in silenced and control mice. These data suggest that during the pre-training phase mice learn a simple association between odor and reward that engages a neural representation of value in the OFC. Once this association is learned, the OFC is no longer required and a second brain structure facilitates the subsequent learning necessary for discrimination.

### Associative Conditioning in Freely Moving Mice

We also examined the role of the OFC during pretraining in a two-phase freely moving behavioral paradigm (Figure 5F). In this task, mice expressing halorhodopsin or YFP in the OFC were placed into an arena. Freely moving mice first learned an association between odor and water during pre-training in an average of 211 trials (Figure 7R, gray), far more trials than required in the head-fixed task. Photoillumination of the OFC severely impaired the ability of 3 of 5 mice expressing halorhodopsin to learn this task (Figure 5G). These mice failed to initiate trials after eight days of training (Figure 5G), whereas the remaining mice initiated trials but learned slower than controls (halorhodopsin: 365 trials, control: 211 trials, p=0.07, ranksum test). The release of inhibition in the OFC restored the ability to learn in 2 of the 3 severely impaired mice. Optrode recordings confirmed that photo-illumination results in a ten-fold inhibition of OFC neurons.

We next examined the consequence of OFC inactivation in the discrimination phase of the task. Freely moving mice expressing either halorhodopsin or YFP in the OFC were pretrained in the absence of inhibition and the OFC was inhibited during discrimination learning. As we observed in the head-fixed task, mice expressing YFP exhibited generalized anticipatory licking at the start of discrimination learning (Figure 7S, 7U, gray). These control mice then suppressed licking to the CS− odor and enhanced licking to the CS+ odor in under 60 trials (Figure 5H, 5I). Licking to CS+ odors (Figure 5H) and suppression of licking for CS− odors (Figure 5I) in silenced mice were similar to controls. The results of OFC inhibition in freely moving mice are in accord with our observations in the head-fixed paradigm and demonstrate that the OFC is important in learning an association between odor and water during pretraining, but once this knowledge is acquired the OFC is no longer necessary for discrimination learning.

### Temporal Representations in OFC in the Two-Phase Paradigm

We performed imaging experiments to examine the relationship between odor representations in OFC and behavior in the two-phase head-fixed task. During pretraining, a strong CS+ representation emerges with 26% of the neurons responding to the pretraining CS+ odor (Figure 6A). Responses after training were more consistent and of higher amplitude than before training and the response power to the CS+ odor increased two-fold (Figure 6E). The properties of this representation are similar to that of the CS+ odors after learning in the single-phase task (Figure 2B). We then performed imaging during discrimination training, with mice exposed to two CS+ and two CS− odors. At the start of discrimination, 21% of the neurons were responsive to each of the four odors (Figure 6B). These responses were non-selective and weak in amplitude (Figure 6B, 6F, 6I). During discrimination training, neurons became selectively responsive to the CS+ odors (Figure 6C). Decoding analysis revealed that the CS+ and CS− ensembles were more separable after discrimination learning (decoder accuracy at onset of discrimination: 0.60, fully learned: 0.93, chance = 0.50) (Figure 6L). However, the response powers to CS+ and CS− odors were significantly weaker than the power to the CS+ odor after pre-training. We continued to image the OFC for up to 4 days after discrimination learning plateaued. The CS+ representation gradually diminished, and the response power decreased to below the response to odors prior to training (Figure 6D, 6H, 6K, S7A). Only 14% of the neurons responded to the CS+ odors and 11% of neurons responded to CS− odors after prolonged training. These imaging results are consistent with the behavioral observations. OFC is required to learn the association of the pre-training odor with water, and a rich representation of this odor is observed upon imaging. OFC is not required for discrimination learning during which its modest CS+ representation weakens considerably. These results suggest that a second structure is employed to accomplish the task of discrimination learning.

**Figure 6.**
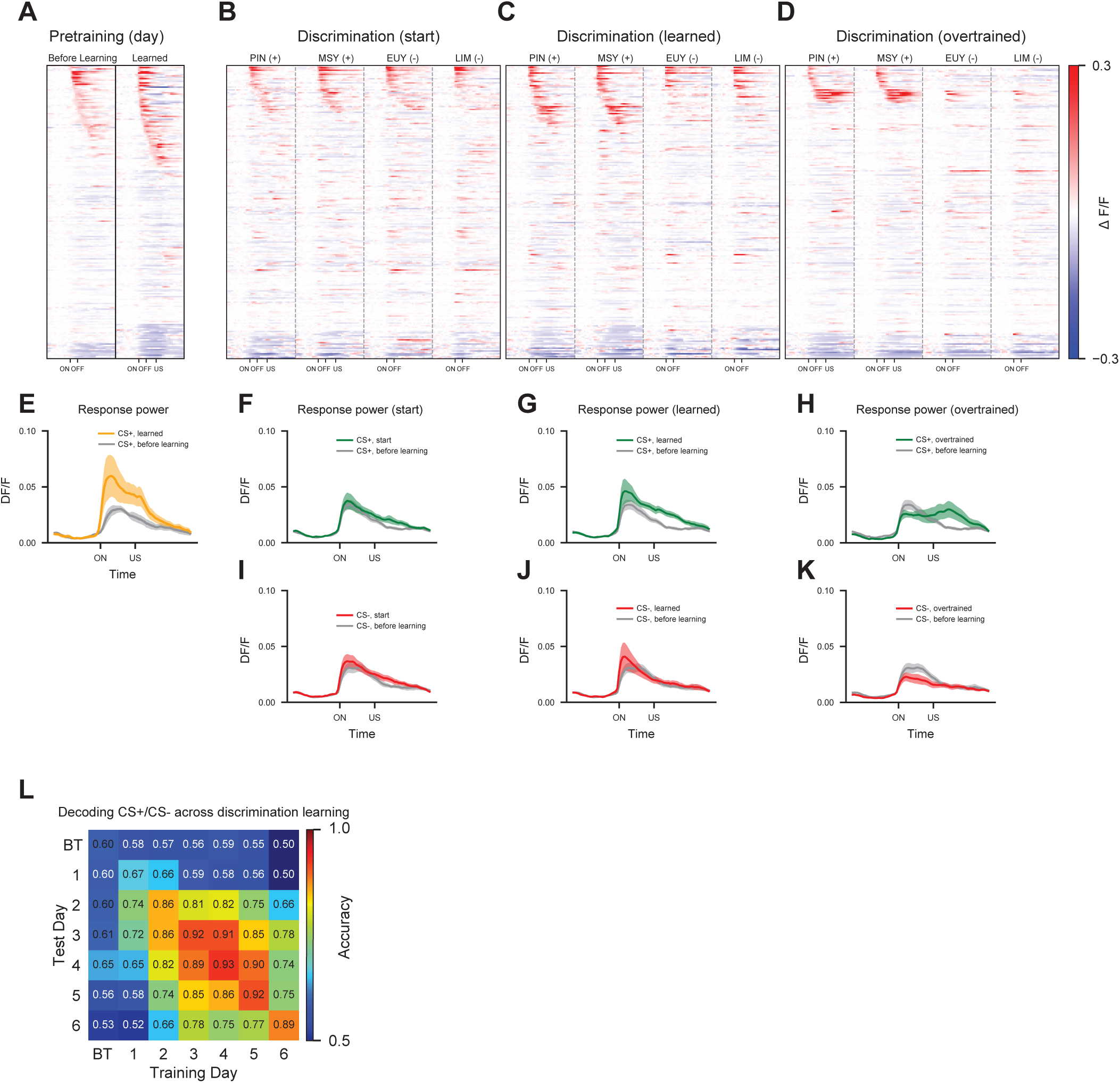
The odor representation in OFC peaks during pretraining and diminishes during discrimination learning. (A-D) PSTH of OFC responses during multiple days of the two-phase task for all mice (n=4). Responses on different days are not aligned but sorted individually. (A) Responses to the CS+ odor (3-octanol) during the pretraining phase before (day 1) and after learning (day 3). (B-D) Responses to 2 new CS+ and 2 CS− odors after the first day of discrimination training (B), after discrimination learning (C), and after over-training (D). (E) Response power of the OFC representation to the CS+ odor before pretraining (gray) and after pretraining (orange). Here and below, shading indicates ±1 SEM. (F-H) CS+ response power on the first day of discrimination learning (F), after discrimination learning (G), and after over-training (H). Average response power evoked by the two CS+ odors during each of these periods (green) is compared to the response power evoked by the same odors prior to training (gray). (I-K) CS− response power on the first day of discrimination learning (I), after discrimination learning (J), and after over-training (K). Average response power evoked by the two CS− odors during each of these periods (red) is compared to the response power evoked by the same odor prior to training (gray). (L) Accuracy of decoding predictive value (CS+ odors vs. CS− odors) from OFC population activity within and across days of discrimination training. 40 randomly chosen neurons per animal were used. Chance accuracy is 50%. BT: before training (naïve odors). Indices for days start after 3-4 days of pretraining has concluded. See also Figure S7.

### A Representation of Value in the mPFC

Previous experiments have implicated the medial prefrontal cortex (mPFC) in reward learning (Birrell and Brown, 2000; Chudasama and Robbins, 2003; Kim et al., 2017; Kitamura et al., 2017; Ostlund and Balleine, 2005; Otis et al., 2017). We therefore performed imaging in the mPFC to discern whether a representation of value emerges during discrimination learning in the two-phase task that may support learning after the diminution of the OFC representation. Imaging of the mPFC during pretraining revealed that the response to the pretraining odor was sparse and of low amplitude, and did not increase with learning (Figure 7A, 7E). Mice were then exposed to two new CS+ and two CS− odors. After one day of discrimination training, we observed a significant response to all odors (Figure 7B, 7F, 7I). The population activities evoked by these odors were more correlated (Figure 7L, 7M) and of higher amplitude (Figure 7F, 7I) than prior to training (Figure S7B), and may reflect the generalized licking to all odors (S6A-D).

**Figure 7.**
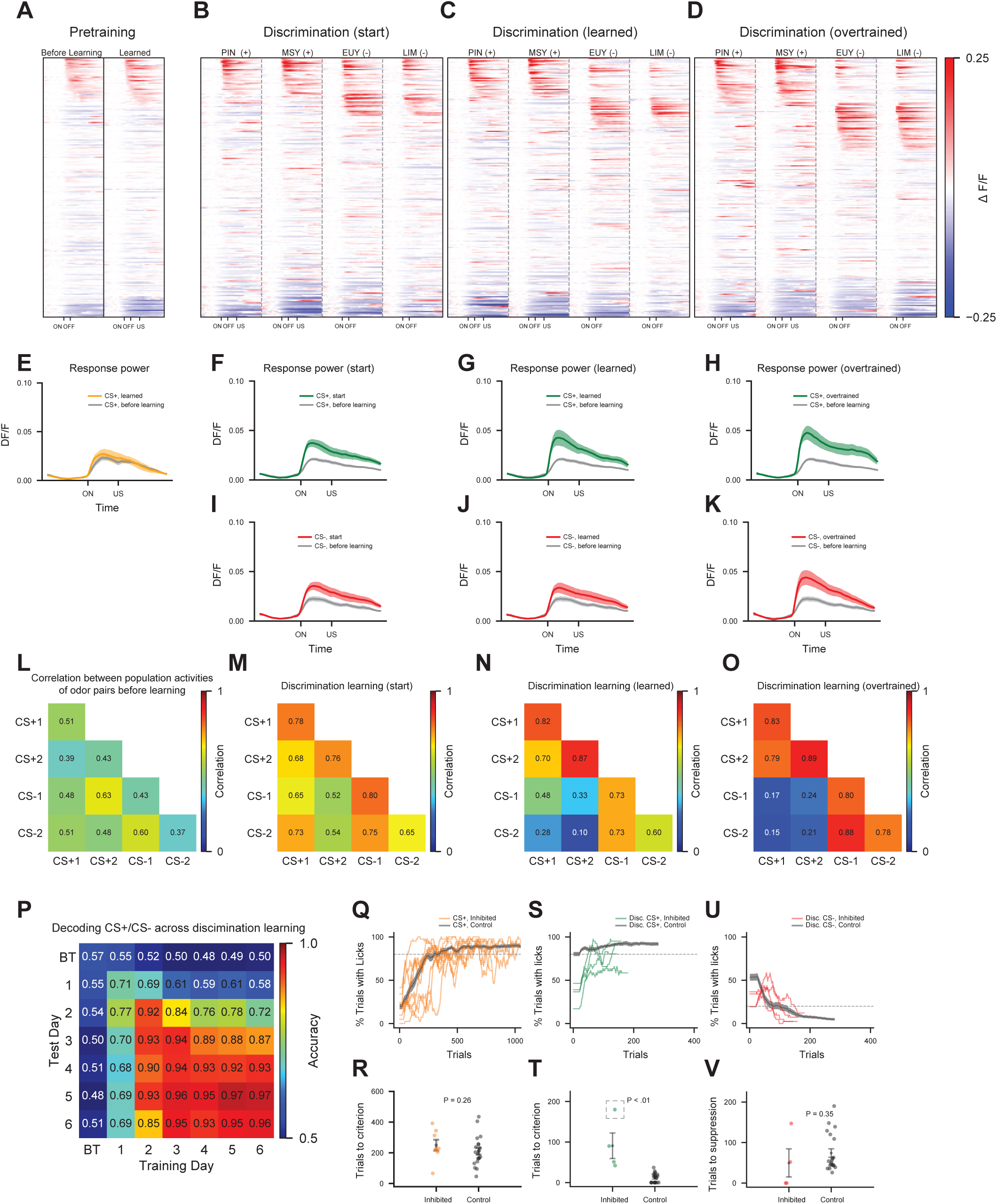
CS+ and CS− representations emerge in mPFC during discrimination and mPFC is required for discrimination learning. (A-D) PSTH of mPFC responses during multiple days of the two-phase task for all mice (n=4). Responses on different days are not aligned but sorted individually. (A) Responses to the CS+ odor during the pretraining phase before and after learning. (B-D) Responses to 2 new CS+ and 2 CS− odors after the first day of discrimination training (B), after discrimination learning (C), and after over-training (D). (E) CS+ response power before pretraining (gray) and after pretraining (orange). Here and below, shading indicates ±1 SEM. (F-H) CS+ response power on the first day of discrimination learning (F), after discrimination learning (G), and after over-training (H). Average response power evoked by the two CS+ odors during each of these periods (green) is compared to the response power evoked by the same odor prior to training (gray). (I-K) CS− response power on the first day of discrimination learning (I), after discrimination learning (J), and after over-training (K). Average response power evoked by the two CS− odors during each of these periods (red) is compared to the response power evoked by the same odor prior to training (gray). (L-O) Within-day correlations between the population activities for all pairs of odors prior to training (L), after the first day of discrimination training (M), after discrimination learning (N), and after over-training (O). Correlation between all pairs of distinct odors before learning (L): 0.52; start of discrimination training (M): 0.64, p = 0.014, Wilcoxon signed-rank test. Correlation between all CS+/CS− odor pairs at start of discrimination training: 0.61, after learning: 0.30, p < 0.001, Wilcoxon signed-rank test. (P) Accuracy of decoding of predictive value (CS+ odors vs. CS− odors) from mPFC population activity within and across days of discrimination training. 40 randomly chosen neurons per animal were used. Chance accuracy is 50%. BT: before training. Indices for days start after 3-4 days of pretraining has concluded. (Q-R) mPFC silencing during the pre-training phase of the two-phase task in freely moving animals. Mice expressing halorhodopsin: n= 8; mice expressing YFP: n= 21. YFP mice are pooled across conditions. (Q) Percentage of trials with anticipatory licking to the pretraining CS+ odor (mice expressing halorhodopsin: orange, mice expressing YFP: gray). Here and below, shading indicates ±1 SEM for control animals. (R) Trials to criterion for licking to the pretraining CS+ odor. Mice expressing halorhodopsin (orange): 249 trials, mice expressing YFP (gray): 211 trials, p = 0.26, ranksum test. Here and below, error bars indicate mean ±1 SEM and dots indicate individual animals. (S-V) mPFC inhibition during the discrimination phase of the two-phase task in freely moving animals. Mice expressing halorhodopsin: n= 4, mice expressing YFP: n= 21. (S) Percentage of trials with anticipatory licking to the CS+ odor (mice expressing halorhodopsin: green, mice expressing YFP: gray). (T) Trials to criterion for licking to the CS+ odor. Mice expressing halorhodopsin (green): 91 trials, mice expressing YFP (gray): 11 trials, p = 0.002, ranksum test. One inhibited mouse did not reach criterion at the end of training (dotted square), and trials to criterion for this mouse was defined as the last trial of training. (U) Percentage of trials with anticipatory licking to the CS− odor (mice expressing halorhodopsin: red, mice expressing YFP: gray). (V) Trials to criterion for suppression of licking to the CS− odor. Mice expressing halorhodopsin (red): 50 trials, mice expressing YFP (gray): 74 trials, p = 0.35, ranksum test. See also Figure S7.

As learning proceeds, we observed a population of neurons responsive only to CS+ odors (19%), accompanied by a second population responsive to CS− odors (22%) (Figure 7C). The CS+ and CS− representations increased in amplitude and became more separable during discrimination learning (Figure 7G, 7J, 7N). We continued to image the mPFC representation for up to 4 days after learning plateaued, and unlike the OFC representations, the mPFC ensembles remained robust and persistent (Figure 7D, 7H, 7K). After prolonged training, 23% of mPFC neurons responded to CS+ odors, and a non-overlapping 25% of mPFC neurons responded to CS− odors (Figure 7D, 7O). These results are further supported by decoding analysis that revealed that the representations of CS+ and CS− odors are stable and separable after discrimination learning (Figure 7P).

We note that whereas we observe a robust CS+ response during discrimination learning, we do not observe a response to the CS+ odor during pretraining. This suggests that the emergence of a CS+ representation in mPFC during discrimination coincide with a requirement to distinguish between CS+ and CS− odors. Whatever the mechanism, the mPFC appears to transform a representation of odor identity encoded in piriform into two distinct and stable representations, a CS+ ensemble encoding positive value and a CS− ensemble encoding negative value.

We next examined the role of mPFC in the two-phase paradigm in freely moving mice. Mice expressing either halorhodopsin or YFP in the mPFC were photoilluminated during the different phases of the task. Inactivation of the mPFC during pretraining did not inhibit task performance (Figure 7Q, 7R), whereas silencing during discrimination impaired appetitive learning in response to CS+ odors. 91 trials on average were required to reach successful learning criterion to CS+ odors in mice (n=4) expressing halorhodopsin compared with 11 trials in control mice (n=21) expressing YFP (Figure 7S, 7T). We note that control mice initiate licking in response to both CS+ and CS− odors at the start of discrimination (Figure 7S, 7U). In contrast, mPFC-silenced mice failed to lick at the onset of discrimination to all odors and slowly learned to lick in response to CS+ odors (Figure 7S, 7T), but not to CS− odors (Figure 7U, 7V). Silencing of the mPFC thus impaired both generalization and discrimination learning, but we do not know whether there is a causal relation between these two components of the behavior.

These data suggest that the neural representation in OFC during pre-training contributes to the learning of an association between odor and water. The OFC representation dissipated upon discrimination learning and a persistent representation of both CS+ and CS− odors emerged in the mPFC. The mPFC supports generalization and participated in the discrimination of odors predictive of reward, suggesting a transfer of information from OFC to mPFC in odor learning.

## DISCUSSION

The representation of odor in piriform cortex is distributive and unstructured and cannot inherently encode value (Illig and Haberly, 2003; Iurilli and Datta, 2017; Poo and Isaacson, 2009; Rennaker et al., 2007; Stettler and Axel, 2009; Sugai et al., 2005; Zhan and Luo, 2010). Value must therefore be imposed in downstream structures by experience or learning. We have examined the representation of odors in piriform cortex as well as in two downstream stations, OFC and mPFC, as mice performed an appetitive olfactory learning task. In piriform cortex we observed minor changes in neural activity unrelated to learning, suggesting that piriform encodes odor identity rather than odor value. In orbitofrontal cortex, 30% of the neurons acquired robust responses to conditioned stimuli after training and these responses were modulated by context and internal state. The representation in OFC, however, dissipated after learning and a more stable representation of both CS+ and CS− odors emerged in medial prefrontal cortex. Thus, the representation of odor identity in piriform is transformed into representations of value in both OFC and mPFC. Moreover, these two brain structures appear to function sequentially in the learning of appetitive associations.

### Representation of Odor Identity in Piriform Cortex

The imposition of value upon neuronal populations is often reflected in changes in either the amplitude or the number of neurons responsive to a stimulus and results in an increase in the sensitivity of the organism to conditioned stimuli (Buonomano and Merzenich, 1998). We observed minor changes in the amplitude and size of neural ensembles responsive to odors during learning in the piriform. Associative learning could also alter the representation of a stimulus in a manner that enhances discrimination (Feldman, 2009). Odor ensembles were slightly more separable upon learning but these changes were qualitatively similar for both CS+ and CS− odors and we6+re also observed upon passive odor exposure. These data suggest that changes in neural populations reflective of value must occur downstream of the piriform cortex.

The imposition of value downstream of piriform may be important to assure the specificity of behavioral output elicited by salient odors (Choi et al., 2011). Imposing value in piriform would result in the modification of outputs to all of piriform’s downstream targets, which could drive multiple behavioral outputs. Value encoded downstream, however, affords a specificity of output that is difficult to achieve if reinforcement is imposed at the level of an odor representation in primary olfactory cortex. In addition, were value imposed in piriform cortex, the gating of value by internal state or external context would limit the perception of odor to subsets of states and contexts. Finally, changes in the weights of either bulbar inputs or associative connections between pyramidal neurons reflective of value could reduce the dimensionality of the odor representation in piriform. Thus, the imposition of value in downstream areas allows the piriform to maintain a high-dimensional representation of odor information that can support flexible and specific associations in multiple downstream regions.

### Representations of Value in OFC and mPFC

We observed that a representation of odor identity in piriform was discarded in the prefrontal cortex and was transformed into a representation of value in OFC and mPFC. After learning, neurons responsive to conditioned stimuli (CS+ odors) emerged in the OFC and these responses were strongly modulated by context and internal state. Distinct representations of CS+ as well as CS− odors subsequently emerged in mPFC. Moreover, silencing of either region of prefrontal cortex impaired different phases of the appetitive learning paradigm. In accord with our findings, a neural representation of rewarded auditory stimuli was identified in both OFC and mPFC, and silencing of these brain structures elicited deficits in the acquisition and expression of learned behavior (Namboodiri et al., 2019; Otis et al., 2017). Representations of conditioned stimuli have been described in multiple brain regions during an associative learning task similar to our behavioral paradigm (Allen et al., 2019; Kim et al., 2017; Namboodiri et al., 2019; Otis et al., 2017). It is perhaps not surprising that prediction of a reward essential for the survival of a thirsty animal would register in multiple brain regions. However, the functional and temporal relationships among the multiple representations and their individual contributions to learning were previously unclear.

We have implemented an associative learning task consisting of two phases, pretraining and discrimination, in an effort to disambiguate the contributions of OFC and mPFC to learning. In the pretraining phase a single odor was paired with water delivery. After mice successfully learned that odor predicts reward, a discrimination phase was initiated in which the animals learned to distinguish CS+ from CS− odors. A representation of the CS+ odor emerged early in OFC during pretraining accompanied by a weaker representation in mPFC. During the discrimination phase the CS+ representation diminished in OFC, whereas representations of CS+ and CS− odors emerged in mPFC and remained stable long after learning. Moreover, silencing of OFC during the pretraining but not the discrimination phase impaired learning, whereas inactivation of mPFC during discrimination but not pretraining impaired learning. These observations suggest that during pretraining mice learn a simple association between odor and reward that engages a representation of value in OFC. Once this association is learned OFC is no longer required for subsequent odor discrimination and the representation in mPFC then facilitates the learning of associations necessary for discrimination.

Previous studies have concluded that lesions of OFC do not impair learning of appetitive associations (Burke et al., 2008; Chudasama and Robbins, 2003; Gallagher et al., 1999; Izquierdo et al., 2004; Ostlund and Balleine, 2007; Schoenbaum et al., 2002; Stalnaker et al., 2007). These studies also employed a two-phase conditioning task, but the consequence of lesioning of OFC during pretraining was not assessed. Similar to our observations, lesioning of OFC did not impair the discrimination phase of learning in these studies. A recent study that used a one-phase task observed that OFC inhibition impairs acquisition of learning, in agreement with our results (Namboodiri et al., 2019). Our studies disambiguate pretraining and discrimination and reveal the importance of OFC in the formation specific of associations between stimulus and reward.

### Representations of Value in Multiple Brain Regions

One interpretation of our findings is that odor learning reinforces piriform inputs to OFC, activating a representation of value. OFC may then teach mPFC during discrimination by reinforcing piriform inputs to this brain structure. In this manner, parallel inputs from piriform to multiple downstream targets can be sequentially reinforced to generate multiple representations of odor value.

There are many potential explanations for the presence of multiple representations of value. Each representation, for example, may have subtly different functions resulting in components of cognitive or behavioral output that are not apparent in simple assays involving licking or freezing. The observation that OFC represents only CS+ responses whereas mPFC encodes both CS+ and CS− responses suggests that the two brain areas have distinct behavioral functions. OFC, for example, may impose value on odors, learning that odor predicts water, whereas odor memory, generalization, and the behavioral distinction between CS+ and CS− odors may require the mPFC.

The observation that the OFC representation precedes that of mPFC may reflect the transfer of information from OFC to mPFC. Contextual fear memory is also thought to require the transfer and consolidation of information. A salient context is initially thought to elicit a representation in CA1 of the hippocampus, which over time reinforces a contextual representation in mPFC (Bontempi et al., 1999; Goshen et al., 2011; Kim and Fanselow, 1992; Kitamura et al., 2017; Squire and Alvarez, 1995; Takehara-Nishiuchi and McNaughton, 2008). At early times after learning, behavior depends on temporal lobe structures. Remote recall, however, depends on mPFC and no longer requires an active hippocampus. The persistence of remote contextual memories after bilateral hippocampal ablations argues for consolidation in cortex dependent upon a reinforcing teaching function mediated by the hippocampus.

Theoretical considerations also reveal advantages to encoding memories in multiple, partitioned brain structures (McClelland et al., 1995; Roxin and Fusi, 2013). The persistence of individual representations depends on the stability of synaptic reinforcement in different brain regions and may dictate their role in the learning process. Plastic synapses effecting fast learning can be rapidly overwritten, whereas less plastic synapses in different brain structures can stabilize memories (Benna and Fusi, 2016; Fusi et al., 2005; Roxin and Fusi, 2013). Whatever the advantage afforded by an early OFC representation, it must be transient because the ensemble of neurons encoding value dissipates while the mPFC representation emerges as a stable ensemble. The transient nature of the OFC population supports models in which OFC performs a teaching function during task acquisition, after which it is no longer required for learning discrimination or for the expression of the learned behavior.

### The OFC Representation is Dependent on State and Context

The ensembles in OFC exhibit features that suggest the incorporation of higher order cognitive information not apparent in the sensory representation in piriform cortex. We observed that both internal state and external context gated the value representation in the OFC. After learning, satiation or alterations in context (removal of the water port) abolished the response to conditioned stimuli. Thus, piriform cortex represents the external world, i.e. the identity of an odor, whereas orbitofrontal and medial prefrontal cortex represent not only the external sensory world, but internal features: learning, context and state. This representation of value in OFC is dependent upon the coincidence of a conditioned stimulus, motivated internal state and appropriate context, and undoubtedly other internal factors that we have not explored. One simple model that incorporates these features invokes direct input of piriform neurons onto pyramidal cells in OFC. OFC neurons may also receive inhibitory inputs that prevent the animal from seeking water when the animal is satiated, and these inhibitory inputs may be disinhibited when the animal is in a thirsty state and in the appropriate context. This model affords a flexibility whereby the same neurons in OFC can represent input from multiple sensory modalities encoding values of distinct valence and gated by different states or contexts.

### Distinct CS+ and CS− Representations in mPFC

Distinct CS+ and CS− representations in mPFC emerged after discrimination learning. The discrimination task we have employed has only two behavioral outcomes: lick or no lick. An explicit representation of each behavioral outcome will likely improve the accuracy of discrimination and may confer flexible responses to changes in stimulus value (Kim et al., 2017). The maintenance of a CS− representation may additionally prevent unlearning when stimulus value changes. Consider a scenario in which a stimulus of positive predictive value no longer predicts reward, and reward is provided once again at a remote time. Experiments on this extinction paradigm reveal that relearning occurred considerably faster than learning a new association with a naïve odor (Bouton, 2004). This suggests that a CS+ representation persists after extinction and provides a stored but silent record of early but extinguished learning. Upon extinction a new CS− representation may emerge for a prior CS+ odor. This CS− representation may be dominant over the persistent CS+ representation to suppress licking. Behavioral extinction will be observed despite the persistence of a positive value representation in mPFC. Thus, the presence of distinct CS+ and CS− representation affords the organism flexible responses in a world of changing value.

### The Generation of Distinct CS+ and CS− Representations

How do representations of CS− and CS+ odors arise in distinct populations of cells in the mPFC? In one model, during pretraining, animals exposed to a single CS+ may learn that odor predicts reward through the emergence of a CS+ representation in the OFC. This CS+ representation in OFC may serve a teaching function in the mPFC at the initiation of discrimination learning and reinforce all piriform inputs onto the mPFC. Early in discrimination learning, all odors will therefore activate the mPFC CS+ ensemble and drive generalized licking behavior. When the animals experience odors that are not associated with reward (CS−) a negative reward prediction error (RPE) signal may be generated by the failure of these odors to predict reward (Schultz, 2016). This negative RPE signal is then relayed onto the mPFC to drive the formation of a CS− ensemble in mPFC, distinct from the CS+ ensemble. In this manner, CS− odors will activate a distinct population of neurons in the mPFC that signals negative value. This model invokes the presence of cognitive representations of odor in at least three different brain regions, each contributing a different component function that ultimately leads to stable, yet flexible memory of stimulus value.

## ACKNOWLEDGMENTS

We thank Cory Root for experimental help with imaging; Abby Zadina, Kristen Lawlor, Yijing Chen, and Matthew Chin for technical assistance; Fabio Stefanini for guidance with decoding; Walter Fischler, Stefano Fusi, Rui Costa, Sandeep Datta, Daniel Salzman, Ashok Litwin-Kumar and members of the Axel and Abbott laboratories for discussions; and Clay Eccard for assistance in the preparation of the manuscript. This work is supported by the Columbia Neurobiology and Behavior Program (P.Y.W.), the Helen Hay Whitney Foundation (C.B.), Simons Foundation (L.F.A and R.A.), NSF NeuroNex Award DBI-1707398 and the Gatsby Charitable Foundation (L.F.A), and the Howard Hughes Medical Institute (R.A.). R.A. is an HHMI investigator.

## AUTHOR CONTRIBUTIONS

P.Y.W, C.B., L.F.A. and R.A. conceived the project, participated in its development and wrote the paper. P.Y.W., C.B., and N.P.S. performed the behavioral experiments and optogenetic manipulations. P.Y.W performed the imaging experiments and data analysis. P.Y.W, W.Z. and P.S. performed optrode experiments.

## DECLARATION OF INTERESTS

The authors declare no competing interests.

## STAR METHODS

### KEY RESOURCES TABLE

| REAGENT OR RESOURCE | SOURCE | IDENTIFIER |
| --- | --- | --- |
| **Experimental Models: Organisms/Strains** |  |  |
| Mouse: wild type C57BL/6J | Jackson Laboratory | 000664 |
| Mouse: rosa26-loxp-stop-loxp-GCaMP6s | Jackson Laboratory | 024106 |
| Mouse: Vglut2-ires-cre | Jackson Laboratory | 016963 |
| **Other** |  |  |
| 200 μm, 0.39 NA optical fiber for optogenetics (mPFC) | Thorlabs | Custom Fabrication |
| 200 μm, 0.39 NA optical fiber for optogenetics (OFC) | Thorlabs | CFM12L02 |
| 0.5-mm GRIN lens | GRINTECH | NEM-050-50-00-920-S-1.5p |
| **Software** |  |  |
| MATLAB | Mathworks | [Mathworks.com](https://www.mathworks.com) |
| MATLAB algorithm for registration within and across imaging sessions | Guizar-Sicairos et al. | https://www.mathworks.com/matlabcentral/fileexchange/18401-efficient-subpixel-image-registration-by-cross-correlation |
| MATLAB algorithm for extracting cellular $CA^{+2}$ signals | Pnevmatikakis et al. | [https://www.cell.com/neuron/fulltext/S0896-6273(15)01084-3](https://www.cell.com/neuron/fulltext/S0896-6273%2815%2901084-3) |
| Custom MATLAB scripts for analyzing $CA^{+2}$ signals | Mathworks | N/A |
| FIJI | University of Wisconsin-Madison LOCI | http://fiji.sc/ |
| Python 3.6 | Python | https://www.python.org/ |
| Scikit-Learn | Pedregosa et al. | https://scikit-learn.org/ |
| iPython and Jupyter | Perez et al. | https://jupyter.org |

### CONTACT FOR REAGENT AND RESOURCE SHARING

Further information and requests for resources and reagents should be directed to and will be fulfilled by the Lead Contact, Richard Axel.

### EXPERIMENTAL MODEL AND SUBJECT DETAILS

All experimental and surgical protocols were performed in accordance with the guide of Care and Use of Laboratory Animals (NIH) and were approved by the Institutional Animal Care and Use Committee at Columbia University. For all head-fixed behavior and inhibition experiments, Vglut2-ires-cre mice (Vong et al., 2011) were crossed to Ai96 (Madisen et al., 2015), and all male and female heterozygous transgenic offspring aged 8-16 weeks were used. For all freely-behaving behavior experiments, C57BL/6J mice aged 8-16 weeks were used. All animals were maintained under a normal 12 hour light/dark cycle with littermates until implantation of optical fibers or GRIN lenses.

### METHOD DETAILS

#### Stereotaxic Surgeries

Mice were anesthetized with ketamine (100 mg/kg) and xylazine (10mg/kg) through intraperitoneal injection and then placed in a stereotactic frame. Body temperature was stabilized using a heating pad attached to a temperature controller. For lens implantation experiments, a 1.0-1.5mm round craniotomy centered on the implantation coordinate was made using a dental drill (see Table S1 for coordinates). Dura and 0.5mm – 1mm of underlying cortex was then aspirated. Blood was washed off at the top constantly through aspiration. A 0.5mm diameter and 6.4 mm length microendoscope was then inserted. After implantation, the microendoscopes were fixed in place using Metabond (Parkell) onto the exposed region. To protect the lens that was protruding out of the skull from damage, a metal enclosure was placed around it (Dytran thread adapter) and covered with an acorn nut (Amazon). Lastly, a custom-made head plate (stainless steel) was attached to the skull with Metabond to allow for head-fixation.

For optical fiber implantation experiments, virus was first injected using a micropipette that was made using a Sutter Micropipette Puller (P-2000). Volumes were injected at 100 nL per minute (see Table S4 for virus and injection information). Afterwards, 0.39-NA optical fibers (Thorlabs) were implanted bilaterally over the desired brain region. Following surgery, mice received buprenorphine (0.05 - 0.1 mg/kg) subcutaneously every 12 hours over the next three days. Mice recovered for at least 4 weeks before the start of any imaging or optogenetic experiment.

#### Animal Behavior

For learning experiments, mice were water-restricted (water bottles taken out of cage) and received water (bottle placed back into cage) for 4-5 minutes every day. Behavioral training began when mice weighed less than 90% of free drinking weight (∼3 days for all experiments). Mice were also weighed every day to ensure good health. No health problems related to dehydration arose at any point.

##### Head-fixed behavior

Mice did not undergo any form of shaping prior to assessment of a learning deficit during either the single-phase head-fixed learning task (Figure 4) or the pre-training phase of the two-phase head-fixed learning task (Figure 5). Mice were head-fixed on a large Styrofoam ball, where they could run freely in one axis (forwards and backwards). During imaging, mouse behavior was monitored with an IR camera (Point Grey). The custom olfactometer was made with mass flow controllers (Aarlborg) and quiet solenoid valves (Lee Company), which are controlled by a USB-DAQ (Measurement Computing) using high voltage transistor arrays. The odor stream was set to 800 mL/min, and split into two equal lines carrying 400 mL / minute (see Table S2 for list of odorants used). One line was dedicated for odor detection by the animal. A narrow opening was placed next to the animal’s nose to allow for odor sampling. The other line was routed to a photo-ionization device (Aurora Scientific) to measure odor ionization, an indicator of odor identity and concentration. Water was delivered through a quiet solenoid-controlled valve (Lee Instruments) to a water port (gavage needle). Licking events were collected through a capacitive touch sensor (Phidgets) attached to the water port. Behavioral training and data acquisition were accomplished with custom MATLAB scripts. All data was collected at 1000 Hz.

Most mice learned instantly, without any prior training, to lick from a water port to collect water. Each odor trial had the following structure: 5 seconds baseline, 2 seconds odor, 3 seconds delay, followed by water in the case of CS+ trials. The inter-trial interval was 25 seconds. During pre-training, only one CS+ odor was presented. In most experiments, octanol served as the CS+ odor during pre-training, and methyl salicylate and pinene served as the CS+ odors, and eucalyptol and limonene served as CS− during discrimination learning. Each day of pre-training consisted of 40-60 trials of the single CS+ odor. Discrimination training consisted of five types of trials, delivered pseudo-randomly: 2 CS+ odors that predicted water delivery, 2 CS− odors, and US trials in which water was delivered without prior odor delivery. Each day of discrimination training consisted of 12-15 trials of each of the 5 conditions (60-75 trials total). For imaging experiments, most training sessions were conducted every other day to minimize GCaMP6S bleaching in transgenic mice.

For bilateral photo-inhibition experiments, a far-red laser (660 nm, CrystaLaser) was used for mice expressing the red-shifted halorhodopsin Jaws, and a 560 nm laser (CrystaLaser) was used for mice expressing the halorhodopsin NpHR. The laser was connected through a single patch cord and a rotary joint (Doric Lenses) to divide the laser output equally onto bilaterally implanted optical fibers. The power at the ends of each fiber tip was approximately 8-10 mW for all inhibition experiments. Laser was turned on 2 seconds prior to odor delivery and turned off 2 seconds after US delivery, lasting for a total of 9 seconds. Laser was also on for 9 seconds in CS− trials, beginning 2 seconds before odor delivery. To confirm that the optical fibers had delivered the expected amount of power during the experiment, fiber implants were extracted immediately after perfusion and output power levels at the fiber tip was re-tested. Additionally, we included only mice in which viral expression within the target area of interest was robust after histological analysis. No mice were excluded based on these two criteria.

##### Freely-moving behavior

Mice did not undergo any form of shaping prior to assessment of a learning deficit during the pre-training phase of the two-phase learning task (Figure 5, 7). Water-restricted animals were placed a in a 1ft x 1ft training chamber and allowed to explore freely. The training chamber was placed in a sound-attenuating PVC cabinet (MedAssociates) and was retrofitted with a custom-made ceiling with a holder (Thorlabs) for a 1 to 2 rotary joint intensity splitter (Doric Lenses) that allowed free movement of the animal during laser photoillumination sessions. The training chamber had a built-in custom-made nose port on one wall. The nose port contained a lick spout (gavage needle) connected to a capacitive touch sensor (Phidgets), a vacuum line connected to wall vacuum and an odor line connected to the olfactometer. Training sessions took place in the dark and animals were monitored with an IR camera (Edmund Optics). All behavioral training was controlled with custom-written Python scripts. Entry of the animals’ nose into the nose port was detected with IR sensors (Sparkfun). All behavioral training was controlled with custom-written Python scripts.

A behavioral training session lasted approximately 30 minutes and an animal could complete as many as 200 trials. For optogenetic silencing experiments, the laser was turned on for the entire training session. The laser output was divided equally to the bilaterally implanted ferrules through the rotary joint. The power was adjusted such that the power coming out of each fiber tip was 10-15 mW for all inhibition experiments. Odors (diluted to 1% with mineral oil) were pinene (CS+ during pre-training), isoamyl acetate (CS+ during discrimination), and ethyl acetate (CS− during discrimination). Odors were delivered with a custom-made olfactometer (mass flow controllers, Aarlborg; quiet solenoid valves, Lee Company; USB-DAQ, National Instruments) and an air pump (MedAssociates) at a rate of 1 L/min. Trials of CS+ and CS− odors were delivered in a pseudo-random order. The trial structure was as follows: the trial was initiated when the animal inserted the nose into the nose port, as detected by the IR sensor. After 0.7 s, if the animal was still in the port (as reported by the IR sensor), the odor was delivered for 2.4 s, followed immediately by water if the odor was a CS+ odor. Each trial was followed by a 5 s inter-trial interval. Behavioral performance was quantified by measuring the percent of time spent licking in the 1.2 s interval before the end of odor delivery.

##### Head-fixed Imaging

A two-photon microscope (Ultima, Bruker) was equipped with the following components to allow imaging of deep brain areas in vivo: a tunable mode-locked 2-photon laser (Chameleon Vision, Coherent) set to 920 nm, ∼100 fs pulse width; a GaAsp-PMT photo-detector with adjustable voltage, gain, and offset feature (Hamamatsu Photonics); a single green/red NDD filter cube (580 dcxd dichroic, hq525/70 m-2p bandpass filter); a long working distance 10X air objective with 0.3 NA (Olympus).

A 260 pixel X 260 pixel region of interest (∼400 um X 400 um FOV) was chosen, with 1.6 us dwell time per pixel, to allow image collection at 4.5 Hz. Imaging from of the same plane across multiple days (z-axis) was accomplished by using the top of the GRIN lens as a reference point for alignment. For each trial, two-photon scanning was triggered at the onset of the baseline period (5 seconds prior to odor delivery), and a 19 second (75 frames) video was collected. Data was acquired using custom acquisition software (Bruker Instruments).

##### Optrode Experiments

Extracellular recordings were performed acutely in head-fixed animals using 32-channel silicon probes (Buzsaki32, NeuroNexus) with a 100 um core fiber attached to one of the four shanks. A 660 nM laser was used for Jaws activation and a 560nm laser for Halorhodopsin activation (CrystaLaser). Recordings were performed 4 weeks after virus injection. On recording days, mice were anesthetized with ketamine/xylazine and the skull indentation created during virus injection was enlarged using a drill and the dura was removed. Subsequently, mice were then head-fixed to the recording stage, and the optrode was lowered inside the brain with a micro-manipulator. The incision was then sealed with liquid agar (1.5%) applied at body temperature.

We lowered optical fibers down to 2-3 mm below Bregma towards the OFC and performed a series of inhibition recordings with varying power levels (.5 mW, 1 mW, 2 mW, 5 mW, 10 mW, and 15 mW) at fiber tip. For each power level, the laser was turned on for 10 seconds with an ITI of 30 seconds for a total of 15 consecutive blocks. In Halorhodopsin-expressing animals, we also performed trials of 10 minutes of photo-illumination to assess OFC silencing in a setting similar to the uninterrupted photo-illumination delivered throughout the entire training session in freely moving experiments.

The 32-channel recording data were digitized at 40 KHz and acquired with OmniPlex D system (Plexon Inc). The voltage signals were high-pass filtered (200 Hz, Bessel) and sorted automatically with KlustaKwik (Rossant et al., 2016) or Kilosort (Pachitariu et al., 2016). The clusters were then manually curated with KlustaViewa or Phy GUI to merge spikes from the same units and to remove instances of noises and units that were not well isolated. Spike data was converted into firing rates using a first-order Savitzky-Golay filter with a smoothing window of 100 ms.

##### Histology

Mice were euthanized after anesthesia with ketamine/xylazine. Brains were extracted and incubated in paraformaldehyde for 24 hours, and then coronal sections (100um) were cut on a vibratome (Leica VT1000 S). The sections were then incubated with far-red neurotrace (640/660, Thermo Fisher Scientific) to label neuronal cell bodies. All images were taken using a Zeiss LSM-710 confocal microscope system. Histology was performed to confirm locations of implanted lenses and optical fibers, as well as expression levels for GCaMP6, YFP, Jaws, and Halorhodopsin.

##### Pooling Animal Cohorts Across Conditions

Animal cohorts A, B, C, and H were pooled as controls for OFC inhibition during the single-phase discrimination learning task (see Tables S3 and S4 for cohort information). Animal cohorts D, E, J, and K were pooled as controls for OFC inhibition during pretraining in the two-phase task. Animal cohorts D, E, and J were pooled as controls for OFC inhibition during discrimination in the two-phase task. Animal cohorts M, O, Q, and S were pooled as controls for inhibition experiments in all freely moving two-phase tasks.

##### Data Collection

Investigators were not blind during either imaging or optogenetic experiments. For imaging experiments, mice were excluded if the field of view contained less than 20 neurons, if the signal was too dim, or if the lens was not placed directly above the region of interest (n=1, Cohort C, Table S3). For optogenetic experiments, mice were excluded if histology revealed low opsin expression within the region of interest, if the optic fibers were not located at the targeted coordinate, or if the optic fibers did not transmit excitation light properly (n=0, all conditions). 1 mouse was excluded from cohort R because it failed to learn pretraining in the two-phase task in the absence of any optogenetic silencing, and we therefore could not assess its performance during subsequent discrimination learning (Table S4).

### QUANTIFICATION AND STATISTICAL ANALYSIS

Image processing and calcium transient analysis were performed using MATLAB. Significance was defined as p < 0.05. All statistical tests, behavioral data analyses and imaging data analyses were performed using Python. Wilcoxon rank-sum test was used in two-group comparisons, Dunn’s test was used for multiple-group comparisons, and Wilcoxon signed-rank test was used in paired group comparisons.

#### Behavioral Data Analysis

For head-fixed behavior, anticipatory licking was defined to be the number of licks within the 1.0 second window prior to water delivery. For freely-moving behavior, anticipatory licking was defined as the percentage of time spent licking in the last 1.2 seconds prior to water delivery. Collection licking was defined as the percentage of time spent licking within the 1.0 seconds after water delivery. AUC (area under ROC) was calculated for each mouse by comparing the distributions of licks to CS+ trials and to CS− trials over a moving window of 20 trials.

Criterion for learning to CS+ odors was defined as the number of trials required to display anticipatory licking in over 80% of the CS+ trials. Criterion for learning to CS− odors was defined as the number of trials required to display any anticipatory licking in less than 20% of the CS− trials. The percentage of trials with anticipatory licking was calculated using a moving window with a length that was adjusted to match the durations to learn in the different tasks. Window lengths: single-phase head-fixed task = 20; pre-training in the two-phase head-fixed task = 20; discrimination in the two-phase head-fixed task = 10; pre-training in the two-phase freely moving task = 40; discrimination in the two-phase discrimination task = 20.

#### Image Processing

Images were first motion-corrected using sub-pixel image registration (Guizar-Sicairos et al., 2008). Motion correction was first applied within each trial (75 frames per trial), and then across trials by registering the mean intensity image of different trials (40-80 trials per imaging session). In some FOVs, we often observed small fluorescence changes occurring in large areas (> 100 um X 100 um) that could be the consequence of calcium transients in out-of-focus planes. We eliminated these diffuse calcium fluctuations through a spatial low-pass Gaussian filter (length constant, 50 um).

#### Calcium Transient Analysis

For ROI identification, we used a MATLAB package for calcium transient analysis based on nonnegative matrix factorization (NMF) (Pnevmatikakis et al., 2016). Spatial filters that corresponded to neurons were selected, and other signals that did not correspond to neural cell bodies (for example, neuropil) were manually deleted. On rare occasions, the algorithm classified distinct neurons in close proximity as one neuron, and the spatial filter was split manually. On average, 70-100 neurons were extracted, and de-noised DF/F was computed for each neuron.

To identify the same neurons across multiple days of imaging sessions, we first performed NMF to extract spatial filters corresponding to neurons on each imaging day. We then aligned image stacks by performing rigid body registration on the mean intensity images (MII) of each imaging day. For example, for a set of imaging data acquired across 5 days, we used the MII on day 3 as reference, and transformation matrices were derived by aligning MIIs on other days relative to day 3. After alignment, we pooled all unique and non-overlapping spatial filters from all imaging days into a single list. Neuronal cell counts obtained after this step typically exceeded standard single-day cell count results by 20-40%. We then back-applied each spatial filter to derive optimal cellular outlines corresponding to the same cell on each imaging day. We manually assessed whether the back-applied spatial filters on each imaging day corresponded to the same cell by evaluating the shapes of the spatial filters while being blind to the fluorescence data. This led to the exclusion of 10-20% of all ROIs when aligning across 4 or more imaging days.

#### Quantification of Significant Neuronal Response

For each cell, we pooled the DF/F values during the baseline period (the first five seconds of each imaging trial prior to odor delivery) of all odor trials to create a reference distribution of DF/F values. This was compared to a distribution of pooled DF/F values centered on a given frame with a moving window of 3 frames (0.67 seconds), in which we refer to as the sample distribution. A Mann-Whitney U test was performed on the reference and sample distributions to obtain a P-value for each frame. Using this method, a P-value was obtained for every frame after odor onset. A cell was defined as significantly active on a given imaging day if: 1) the P-value was less than 0.01 for at least 8 consecutive frames after odor onset and 2) the maximum DF/F during the odor delivery period exceeded the DF/F during the baseline period over a set threshold. This DF/F threshold was 0.10 for piriform responses, 0.04 for OFC responses, and 0.03 for mPFC responses to account for observed differences in GCaMP6s expression within each area in our transgenic mice. We used this metric to quantify the fraction of cells responsive to a stimulus on a given imaging day.

After reversal learning, OFC responses to the old CS+ odors in most neurons did not diminish down to amplitudes observed prior to training. We thus quantified the fraction of neurons that responded more to CS+ odors than to CS− odors, and vice versa, after discrimination learning and after reversal learning. For a neuron to be considered to be responding more to CS+ odors than CS− odors, it must have statistically significant responses (see above) to both CS+ odors and also respond with greater amplitude to CS+ odors than to CS− odors.

#### Response Power

We defined the response power of a neuronal population to a given odor to be the mean excitatory response to that odor. Neuronal responses that were inhibitory throughout the entire odor presentation period and the delay period were excluded from this analysis because the magnitude of inhibition cannot be reliably measured with genetically encoded calcium indicators, especially given low baseline firing rates.

#### Correlation Analysis

The maximum trial-averaged DF/F response between odor onset and water onset for all neurons was computed, forming a vector that corresponds to the population activity evoked by a given odor. The Pearson product-moment correlation coefficient was then calculated based on two such population activity vectors. This was used to compute the correlation of population activities evoked by the same odor across different training days as well as the correlation of population activities evoked by different odors within a training day.

To correlate an odor ensemble with itself (for example, the diagonal entries of Figure 1H), trials of a given odor were split into two equal halves after random shuffling, and the correlation was then calculated using the trial-averaged DF/F responses of the split data. This was repeated 100 times.

#### Decoding

Support vector machines with linear kernels were constructed using the scikit-learn library in Python. For each odor trial, we created a vector that corresponds to the population activity, based on the maximum DF/F between odor onset and water onset for each trial. The number of neurons used was standardized across all animals in all conditions to be 40 and they were randomly chosen. For the decoding of odors across days, we trained the decoder using odor trials from a given day and tested decoding performance with odor trials on all other days. For decoding trials within the same day, we trained decoders using 5-fold cross-validation. Decoding simulations were repeated 100 times per condition, drawing a new and random set of 40 neurons for each condition.

For decoding odor value, CS+1 and CS+2 odor trials were pooled together, and CS-1 and CS-2 were pooled together, and each group had a different label. For decoding CS+ odor identity, only CS+1 trials and CS+2 trials were used, and each group had a different label. For decoding CS− odor identity, only CS-1 trials and CS-2 trials were used, and each group had a different label. For decoding odor identity, CS+1, CS+2, CS-1, and CS-2 trials were used and have different labels. The strategy used for multi-class decoding of odor identities is the “one-against-one” multi-class classification approach. The values for chance performance for these conditions using random shuffling with 50 repetitions were: odor valence 50%, CS+ identity 50%, CS− identity 50%, odor identity 25%.

## References

Allen, W.E., Chen, M.Z., Pichamoorthy, N., Tien, R.H., Pachitariu, M., Luo, L., and Deisseroth, K. (2019). Thirst regulates motivated behavior through modulation of brainwide neural population dynamics. Science (80-.).

Barretto, R.P., Messerschmidt, B., and Schnitzer, M.J. (2009). In vivo fluorescence imaging with high-resolution microlenses. Nat Methods 6, 511–512.

Benna, M.K., and Fusi, S. (2016). Computational principles of synaptic memory consolidation. Nat. Neurosci.

Birrell, J.M., and Brown, V.J. (2000). Medial Frontal Cortex Mediates Perceptual Attentional Set Shifting in the Rat. J. Neurosci.

Bissonette, G.B., Martins, G.J., Franz, T.M., Harper, E.S., Schoenbaum, G., and Powell, E.M. (2008). Double Dissociation of the Effects of Medial and Orbital Prefrontal Cortical Lesions on Attentional and Affective Shifts in Mice. J. Neurosci.

Bontempi, B., Laurent-Demir, C., Destrade, C., and Jaffard, R. (1999). Time-dependent reorganization of brain circuitry underlying long-term memory storage. Nature 400, 671–675.

Bouton, M.E. (2004). Context and behavioral processes in extinction. Learn. Mem.

Bozza, T., McGann, J.P., Mombaerts, P., and Wachowiak, M. (2004). In vivo imaging of neuronal activity by targeted expression of a genetically encoded probe in the mouse. Neuron 42, 9–21.

Buck, L., and Axel, R. (1991). A novel multigene family may encode odorant receptors: a molecular basis for odor recognition. Cell 65, 175–187.

Buonomano, D. V, and Merzenich, M.M. (1998). Cortical plasticity: from synapses to maps. Annu. Rev. Neurosci. 21, 149–186.

Burke, K.A., Franz, T.M., Miller, D.N., and Schoenbaum, G. (2008). The role of the orbitofrontal cortex in the pursuit of happiness and more specific rewards. Nature.

Chen, C.F., Zou, D.J., Altomare, C.G., Xu, L., Greer, C.A., and Firestein, S.J. (2014). Nonsensory target-dependent organization of piriform cortex. Proc Natl Acad Sci U S A 111, 16931–16936.

Chen, T.W., Wardill, T.J., Sun, Y., Pulver, S.R., Renninger, S.L., Baohan, A., Schreiter, E.R., Kerr, R.A., Orger, M.B., Jayaraman, V., et al. (2013). Ultrasensitive fluorescent proteins for imaging neuronal activity. Nature 499, 295–300.

Choi, G.B., Stettler, D.D., Kallman, B.R., Bhaskar, S.T., Fleischmann, A., and Axel, R. (2011). Driving opposing behaviors with ensembles of piriform neurons. Cell 146, 1004–1015.

Chudasama, Y., and Robbins, T.W. (2003). Dissociable contributions of the orbitofrontal and infralimbic cortex to pavlovian autoshaping and discrimination reversal learning: further evidence for the functional heterogeneity of the rodent frontal cortex. J. Neurosci. 23, 8771–8780.

Chuong, A.S., Miri, M.L., Busskamp, V., Matthews, G.A.C., Acker, L.C., Sørensen, A.T., Young, A., Klapoetke, N.C., Henninger, M.A., Kodandaramaiah, S.B., et al. (2014). Noninvasive optical inhibition with a red-shifted microbial rhodopsin. Nat. Neurosci. 17, 1123–1129.

Critchley, H.D., and Rolls, E.T. (1996). Hunger and satiety modify the responses of olfactory and visual neurons in the primate orbitofrontal cortex. J Neurophysiol 75, 1673–1686.

Davison, I.G., and Ehlers, M.D. (2011). Neural circuit mechanisms for pattern detection and feature combination in olfactory cortex. Neuron 70, 82–94.

Denk, W., Strickler, J.H., and Webb, W.W. (1990). Two-photon laser scanning fluorescence microscopy. Science (80-.).

Diodato, A., Ruinart De Brimont, M., Yim, Y.S., Derian, N., Perrin, S., Pouch, J., Klatzmann, D., Garel, S., Choi, G.B., and Fleischmann, A. (2016). Molecular signatures of neural connectivity in the olfactory cortex. Nat. Commun.

Feierstein, C.E., Quirk, M.C., Uchida, N., Sosulski, D.L., and Mainen, Z.F. (2006). Representation of spatial goals in rat orbitofrontal cortex. Neuron 51, 495–507.

Feldman, D.E. (2009). Synaptic Mechanisms for Plasticity in Neocortex. Annu. Rev. Neurosci.

Ferenczi, E.A., Zalocusky, K.A., Liston, C., Grosenick, L., Warden, M.R., Amatya, D., Katovich, K., Mehta, H., Patenaude, B., Ramakrishnan, C., et al. (2016). Prefrontal cortical regulation of brainwide circuit dynamics and reward-related behavior. Science (80-.).

Fusi, S., Drew, P.J., and Abbott, L.F. (2005). Cascade models of synaptically stored memories. Neuron.

Gallagher, M., McMahan, R.W., and Schoenbaum, G. (1999). Orbitofrontal cortex and representation of incentive value in associative learning. J Neurosci 19, 6610–6614.

Ghosh, S., Larson, S.D., Hefzi, H., Marnoy, Z., Cutforth, T., Dokka, K., and Baldwin, K.K. (2011). Sensory maps in the olfactory cortex defined by long-range viral tracing of single neurons. Nature 472, 217–220.

Godfrey, P.A., Malnic, B., and Buck, L.B. (2004). The mouse olfactory receptor gene family. Proc. Natl. Acad. Sci. 101, 2156–2161.

Goshen, I., Brodsky, M., Prakash, R., Wallace, J., Gradinaru, V., Ramakrishnan, C., and Deisseroth, K. (2011). Dynamics of Retrieval Strategies for Remote Memories. Cell 147, 678–689.

Gottfried, J.A., O’Doherty, J., and Dolan, R.J. (2003). Encoding predictive reward value in human amygdala and orbitofrontal cortex. Science 301, 1104–1107.

Gradinaru, V., Thompson, K.R., and Deisseroth, K. (2008). eNpHR: a Natronomonas halorhodopsin enhanced for optogenetic applications. Brain Cell Biol 36, 129–139.

Guizar-Sicairos, M., Thurman, S.T., and Fienup, J.R. (2008). Efficient subpixel image registration algorithms. Opt. Lett.

Illig, K.R., and Haberly, L.B. (2003). Odor-evoked activity is spatially distributed in piriform cortex. J. Comp. Neurol.

Iurilli, G., and Datta, S.R. (2017). Population Coding in an Innately Relevant Olfactory Area. Neuron.

Izquierdo, A., Suda, R.K., and Murray, E.A. (2004). Bilateral orbital prefrontal cortex lesions in rhesus monkeys disrupt choices guided by both reward value and reward contingency. J. Neurosci. 24, 7540–7548.

Johnson, D.M., Illig, K.R., Behan, M., and Haberly, L.B. (2000). New features of connectivity in piriform cortex visualized by intracellular injection of pyramidal cells suggest that “primary” olfactory cortex functions like “association” cortex in other sensory systems. J. Neurosci. 20, 6974–6982.

Jung, J.C., Mehta, A.D., Aksay, E., Stepnoski, R., and Schnitzer, M.J. (2004). In Vivo Mammalian Brain Imaging Using One- and Two-Photon Fluorescence Microendoscopy. J. Neurophysiol.

Kepecs, A., Uchida, N., Zariwala, H.A., and Mainen, Z.F. (2008). Neural correlates, computation and behavioural impact of decision confidence. Nature 455, 227–231.

Kim, J.J., and Fanselow, M.S. (1992). Modality-specific retrograde amnesia of fear. Science (80-.).

Kim, C.K., Ye, L., Jennings, J.H., Pichamoorthy, N., Tang, D.D., Yoo, A.C.W., Ramakrishnan, C., and Deisseroth, K. (2017). Molecular and Circuit-Dynamical Identification of Top-Down Neural Mechanisms for Restraint of Reward Seeking. Cell.

Kitamura, T., Ogawa, S.K., Roy, D.S., Okuyama, T., Morrissey, M.D., Smith, L.M., Redondo, R.L., and Tonegawa, S. (2017). Engrams and circuits crucial for systems consolidation of a memory. Science 356, 73–78.

Lipton, P.A., Alvarez, P., and Eichenbaum, H. (1999). Crossmodal associative memory representations in rodent orbitofrontal cortex. Neuron 22, 349–359.

Madisen, L., Garner, A.R., Shimaoka, D., Chuong, A.S., Klapoetke, N.C., Li, L., van der Bourg, A., Niino, Y., Egolf, L., Monetti, C., et al. (2015). Transgenic mice for intersectional targeting of neural sensors and effectors with high specificity and performance. Neuron 85, 942–958.

McClelland, J.L., McNaughton, B.L., and O’Reilly, R.C. (1995). Why there are complementary learning systems in the hippocampus and neocortex: Insights from the successes and failures of connectionist models of learning and memory. Psychol. Rev.

Miyamichi, K., Amat, F., Moussavi, F., Wang, C., Wickersham, I., Wall, N.R., Taniguchi, H., Tasic, B., Huang, Z.J., He, Z., et al. (2011). Cortical representations of olfactory input by trans-synaptic tracing. Nature 472, 191–196.

Mombaerts, P., Wang, F., Dulac, C., Chao, S.K., Nemes, A., Mendelsohn, M., Edmondson, J., and Axel, R. (1996). Visualizing an Olfactory Sensory Map. Cell 87, 675–686.

Namboodiri, V.M.K., Otis, J.M., van Heeswijk, K., Voets, E.S., Alghorazi, R.A., Rodriguez-Romaguera, J., Mihalas, S., and Stuber, G.D. (2019). Single-cell activity tracking reveals that orbitofrontal neurons acquire and maintain a long-term memory to guide behavioral adaptation. Nat. Neurosci.

Ostlund, S.B., and Balleine, B.W. (2005). Lesions of medial prefrontal cortex disrupt the acquisition but not the expression of goal-directed learning. J. Neurosci. 25, 7763–7770.

Ostlund, S.B., and Balleine, B.W. (2007). Orbitofrontal Cortex Mediates Outcome Encoding in Pavlovian But Not Instrumental Conditioning. J. Neurosci.

Otis, J.M., Namboodiri, V.M.K., Matan, A.M., Voets, E.S., Mohorn, E.P., Kosyk, O., McHenry, J.A., Robinson, J.E., Resendez, S.L., Rossi, M.A., et al. (2017). Prefrontal cortex output circuits guide reward seeking through divergent cue encoding. Nature 543, 103–107.

Pachitariu, M., Steinmetz, N.A., Kadir, S.N., Carandini, M., and Harris, K.D. (2016). Fast and accurate spike sorting of high-channel count probes with KiloSort. In Advances in Neural Information Processing Systems 29, D.D. Lee, M. Sugiyama, U. V Luxburg, I. Guyon, and R. Garnett, eds. (Curran Associates, Inc.), pp. 4448–4456.

Padoa-Schioppa, C., and Assad, J.A. (2006). Neurons in the orbitofrontal cortex encode economic value. Nature 441, 223–226.

Pnevmatikakis, E.A., Soudry, D., Gao, Y., Machado, T.A., Merel, J., Pfau, D., Reardon, T., Mu, Y., Lacefield, C., Yang, W., et al. (2016). Simultaneous Denoising, Deconvolution, and Demixing of Calcium Imaging Data. Neuron 89, 285–299.

Poo, C., and Isaacson, J.S. (2009). Odor representations in olfactory cortex: “sparse” coding, global inhibition, and oscillations. Neuron 62, 850–861.

Price, J.L. (1985). Beyond the primary olfactory cortex: Olfactory-related areas in the neocortex, thalamus and hypothalamus. Chem. Senses.

Price, J.L., and Powell, T.P. (1970). The mitral and short axon cells of the olfactory bulb. J Cell Sci 7, 631–651.

Ramus, S.J., and Eichenbaum, H. (2000). Neural correlates of olfactory recognition memory in the rat orbitofrontal cortex. J Neurosci 20, 8199–8208.

Rennaker, R.L., Chen, C.F., Ruyle, A.M., Sloan, A.M., and Wilson, D.A. (2007). Spatial and temporal distribution of odorant-evoked activity in the piriform cortex. J Neurosci 27, 1534–1542.

Ressler, K.J., Sullivan, S.L., and Buck, L.B. (1993). A zonal organization of odorant receptor gene expression in the olfactory epithelium. Cell 73, 597–609.

Ressler, K.J., Sullivan, S.L., and Buck, L.B. (1994). Information coding in the olfactory system: evidence for a stereotyped and highly organized epitope map in the olfactory bulb. Cell 79, 1245–1255.

Roesch, M.R., Stalnaker, T.A., and Schoenbaum, G. (2007). Associative encoding in anterior piriform cortex versus orbitofrontal cortex during odor discrimination and reversal learning. Cereb. Cortex 17, 643–652.

Rossant, C., Kadir, S.N., Goodman, D.F.M., Schulman, J., Hunter, M.L.D., Saleem, A.B., Grosmark, A., Belluscio, M., Denfield, G.H., Ecker, A.S., et al. (2016). Spike sorting for large, dense electrode arrays. Nat. Neurosci.

Roxin, A., and Fusi, S. (2013). Efficient Partitioning of Memory Systems and Its Importance for Memory Consolidation. PLoS Comput. Biol.

Royer, S., Zemelman, B. V., Barbic, M., Losonczy, A., Buzsáki, G., and Magee, J.C. (2010). Multi-array silicon probes with integrated optical fibers: Light-assisted perturbation and recording of local neural circuits in the behaving animal. Eur. J. Neurosci.

Schoenbaum, G., and Eichenbaum, H. (1995). Information Coding in the Rodent Prefrontal Cortex .1. Single-Neuron Activity in Orbitofrontal Cortex Compared with That in Pyriform Cortex. J. Neurophysiol. 74, 733–750.

Schoenbaum, G., Chiba, A.A., and Gallagher, M. (1998). Orbitofrontal cortex and basolateral amygdala encode expected outcomes during learning. Nat. Neurosci. 1, 155–159.

Schoenbaum, G., Chiba, A.A., and Gallagher, M. (1999). Neural encoding in orbitofrontal cortex and basolateral amygdala during olfactory discrimination learning. J. Neurosci. 19, 1876–1884.

Schoenbaum, G., Nugent, S.L., Saddoris, M.P., and Setlow, B. (2002). Orbitofrontal lesions in rats impair reversal but not acquisition of go, no-go odor discriminations. Neuroreport 13, 885–890.

Schoenbaum, G., Setlow, B., Nugent, S.L., Saddoris, M.P., and Gallagher, M. (2003). Lesions of orbitofrontal cortex and basolateral amygdala complex disrupt acquisition of odor-guided discriminations and reversals. Learn Mem 10, 129–140.

Schultz, W. (2016). Dopamine reward prediction error coding. Dialogues Clin. Neurosci.

Schwabe, K., Ebert, U., and Loscher, W. (2004). The central piriform cortex: anatomical connections and anticonvulsant effect of GABA elevation in the kindling model. Neuroscience 126, 727–741.

Sosulski, D.L., Bloom, M.L., Cutforth, T., Axel, R., and Datta, S.R. (2011). Distinct representations of olfactory information in different cortical centres. Nature 472, 213–216.

Squire, L.R., and Alvarez, P. (1995). Retrograde amnesia and memory consolidation: a neurobiological perspective. Curr. Opin. Neurobiol.

Stalnaker, T.A., Franz, T.M., Singh, T., and Schoenbaum, G. (2007). Basolateral Amygdala Lesions Abolish Orbitofrontal-Dependent Reversal Impairments. Neuron.

Stettler, D.D., and Axel, R. (2009). Representations of Odor in the Piriform Cortex. Neuron 63, 854–864.

Sugai, T., Miyazawa, T., Fukuda, M., Yoshimura, H., and Onoda, N. (2005). Odor-concentration coding in the guinea-pig piriform cortex. Neuroscience.

Takehara-Nishiuchi, K., and McNaughton, B.L. (2008). Spontaneous changes of neocortical code for associative memory during consolidation. Science (80-.).

Thorpe, S.J., Rolls, E.T., and Maddison, S. (1983). The orbitofrontal cortex: neuronal activity in the behaving monkey. Exp Brain Res 49, 93–115.

Tremblay, L., and Schultz, W. (1999). Relative reward preference in primate orbitofrontal cortex. Nature 398, 704–708.

Vassar, R., Chao, S.K., Sitcheran, R., Nuñez, J.M., Vosshall, L.B., and Axel, R. (1994). Topographic organization of sensory projections to the olfactory bulb. Cell 79, 981–991.

Vong, L., Ye, C., Yang, Z., Choi, B., Chua Jr., S., and Lowell, B.B. (2011). Leptin action on GABAergic neurons prevents obesity and reduces inhibitory tone to POMC neurons. Neuron 71, 142–154.

Zhan, C., and Luo, M. (2010). Diverse patterns of odor representation by neurons in the anterior piriform cortex of awake mice. J Neurosci 30, 16662–16672.

Zhang, X., and Firestein, S. (2002). The olfactory receptor gene superfamily of the mouse. Nat Neurosci 5, 124–133.

