## Supplemental data for "The Imposition of Value on Odor: Transient and Persistent Representations of Odor Value in Prefrontal Cortex"

### SUPPLEMENTAL INFORMATION

#### SUPPLEMENTARY FIGURES

Figure S1

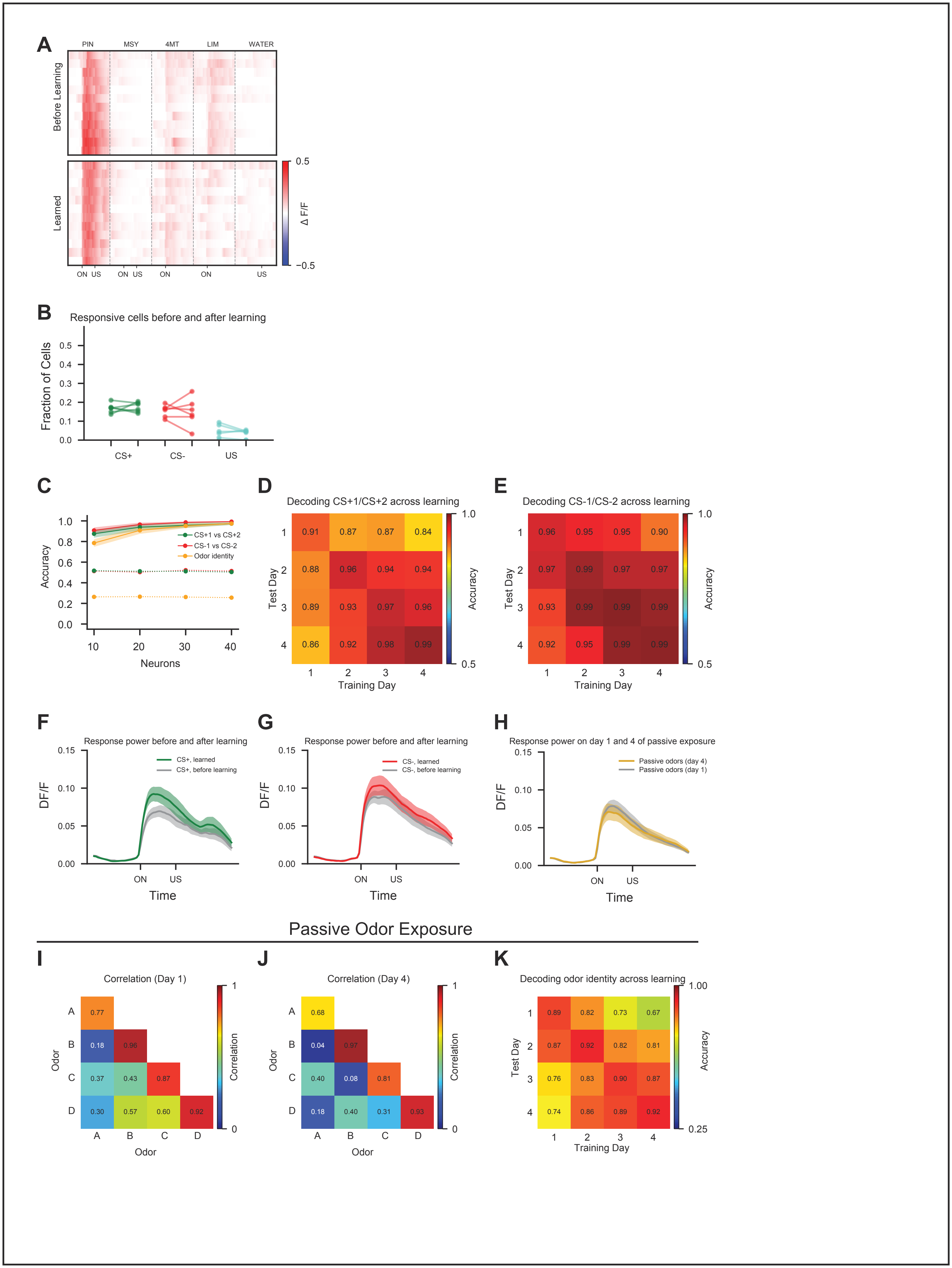
A
PIN
MSY
4MT
LIM
WATER
Before Learning
Learned
0.5
Δ F/F
−0.5
ON US
ON US
ON
ON
US
B
Responsive cells before and after learning
Fraction of Cells
CS+
CS-
US
C
Accuracy
CS+1 vs CS+2
CS-1 vs CS-2
Odor identity
Neurons
D
Decoding CS+1/CS+2 across learning
Test Day
Training Day
Accuracy
E
Decoding CS-1/CS-2 across learning
F
Response power before and after learning
CS+, learned
CS+, before learning
DF/F
Time
G
Response power before and after learning
CS-, learned
CS-, before learning
H
Response power on day 1 and 4 of passive exposure
Passive odors (day 4)
Passive odors (day 1)
Passive Odor Exposure
I
Correlation (Day 1)
Odor
Correlation
J
Correlation (Day 4)
K
Decoding odor identity across learning

**Figure S1. Analysis of piriform responses during learning and passive odor exposure. Related to Figure 1**

(A) Odor responses of an example piriform neuron (1$^{st}$ cell in Figure 1D) on individual trials before learning (top panel, day 1 of training) and after learning (bottom panel, day 4 of training). ON: odor onset. US: water delivery.

(B) Fraction of cells responsive to CS+ odors, CS- odors, and water before and after learning (n = 6 mice). Responses to the two CS+ odors and to the two CS- odors are averaged for each mouse. Dots represent individual animals. Average fraction responsive to CS+ odors before learning: 0.17, after learning: 0.17, $p > 0.05$; CS- odors before learning: 0.15, after learning: 0.15, $p > 0.05$; US before learning: 0.04, after learning: 0.03, $p > 0.05$. Wilcoxon signed-rank test. See STAR Methods for quantification of responsive cells.

(C) Accuracy of decoding the identities of the 4 training odors (orange), CS+1 vs. CS+2 (green), and CS-1 vs. CS-2 (red) from piriform population activity after learning, plotted as a function of the number of neurons used in the decoder. Dotted lines represent chance levels for each experimental condition. Here and below, shading indicates ±1 SEM.

(D and E) Accuracy of decoding CS+1 vs CS+2 (D) and CS-1 vs CS-2 (E) from population activity within and across training days, as in Figure 1J. CS+1/CS+2 before learning (day 1): 0.91, after learning (day 4): 0.99, $p = 0.02$. CS-1/CS-2 before learning (day 1): 0.96, after learning (day 4): 0.99, $p > 0.05$. Wilcoxon signed-rank test.

(F-G) Average response power of piriform neurons to CS+ (F) and CS- (G) odors before learning (gray), and after learning (CS+: green, CS-: red). See STAR Methods.

(H) Average response power of piriform neurons to odors on day 1 (gray) and day 4 of passive odor exposure (orange). n=4 mice.

(I and J) Correlation of activity evoked by all odor pairs on day 1 (I) and day 4 (J) of passive odor exposure ($n = 4$ mice). Correlation for all pairs of distinct odors before learning: 0.41, after learning: 0.23, $p < 0.001$, Wilcoxon signed-rank test. Odors are A: 2-phenylethanol, B: benzaldehyde, C: methyl salicylate, D: octanol.

(K) Accuracy of decoding the identities of the four tested odors from piriform population activity within and across days of passive odor exposure. CS-1/CS-2 before learning (day 1): 0.89, after learning (day 4): 0.92, $p > 0.05$.

### Figure S2

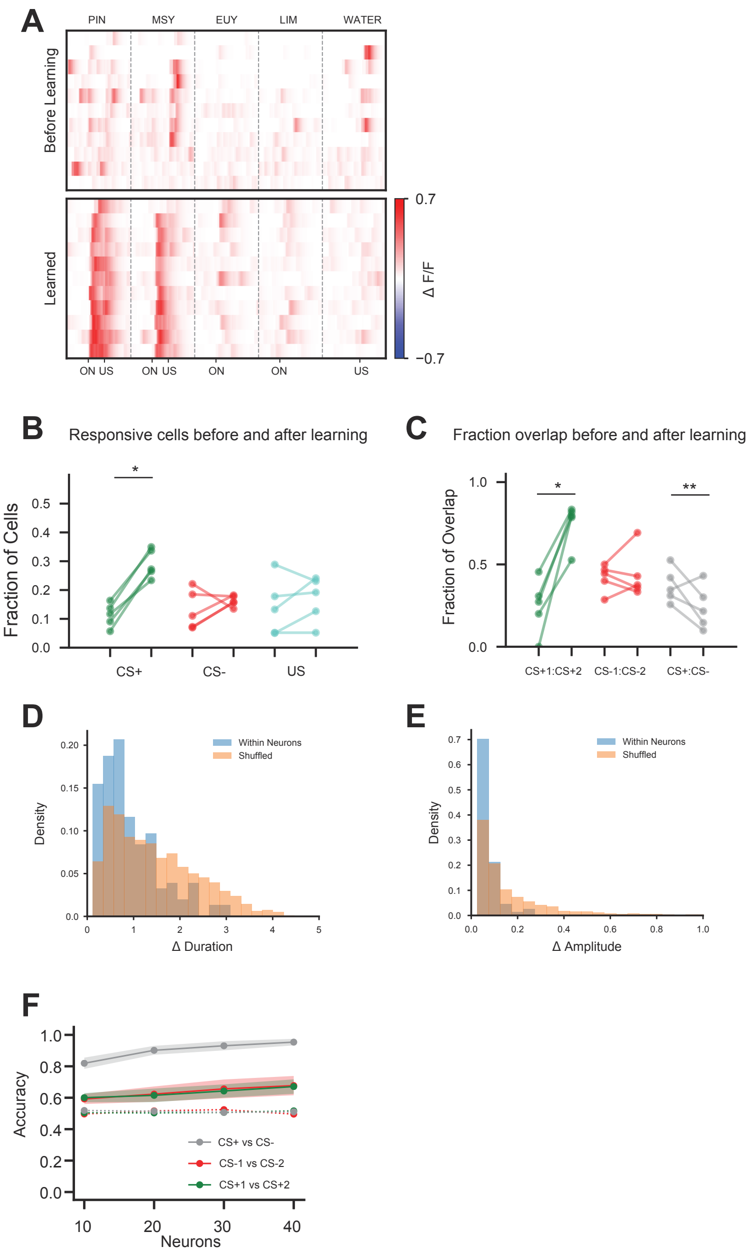

A
PIN
MSY
EUY
LIM
WATER
Before Learning
Learned
ON US
ON US
ON
ON
US
0.7
Δ F/F
−0.7
B
Responsive cells before and after learning
Fraction of Cells
CS+
CS-
US
C
Fraction overlap before and after learning
Fraction of Overlap
CS+1:CS+2
CS-1:CS-2
CS+:CS-
D
Within Neurons
Shuffled
Density
Δ Duration
E
Within Neurons
Shuffled
Density
Δ Amplitude
F
Accuracy
CS+ vs CS-
CS-1 vs CS-2
CS+1 vs CS+2
Neurons

**Figure S2. Analysis of OFC responses before and after learning. Related to Figure 2**

(A) Odor responses of an example OFC neuron (3rd cell in Figure 2C) on individual trials before learning (top panel, day 1 of training) and after learning (bottom panel, day 4 of training). ON: odor onset. US: water delivery.

(B) Fraction of cells responsive to CS+ odors, CS- odors, and water before and after learning (n = 5 mice). Average fraction responsive to CS+ odors before learning: 0.11, after learning: 0.30, p = 0.04; CS- odors before learning: 0.13, after learning: 0.16, p > 0.05; US before learning: 0.14, after learning: 0.17, p > 0.05. Wilcoxon signed-rank test. See STAR Methods for quantification of responsive cells. Here and in C, dots represent individual animals.

(C) Fraction of overlap between cells responsive to CS+1 and CS+2 (green), to CS-1 and CS-2 (red), and to all CS+/CS- odor pairs (gray) before and after learning. Overlap was calculated as $(A \cap B) / \max(A, B)$. Average fraction overlap between CS+1/CS+2 before learning: 0.25, after learning: 0.75, p = 0.04; CS-1/CS-2 before learning: 0.42, after learning: 0.44, p > 0.05; CS+/CS- before learning: 0.37, after learning: 0.24, p = 0.005. Wilcoxon signed-rank test.

(D and E) Similarity of response amplitude (D) and response duration (E) to CS+1 and CS+2 in CS+ responsive neurons. The difference in the amplitude and duration of responses to CS+1 and CS+2 within each neuron (blue) is compared to the difference across different neurons after 100 iterations of shuffling (orange). The distribution of differences is plotted for all CS+ responsive neurons after learning (n=5 mice). The response amplitude and duration evoked by the CS+ odors within each neuron are more similar than across neurons ($p < 0.001$ for both duration and amplitude, ranksum test).

(F) Accuracy of decoding CS+ vs. CS- (gray), CS+1 vs. CS+2 (green), and CS-1 vs. CS-2 (red) from OFC population activity after learning, plotted as a function of the number of neurons used in the decoder. Shading indicates ±1 SEM. Dotted lines represent chance levels for each experimental condition.

### Figure S3

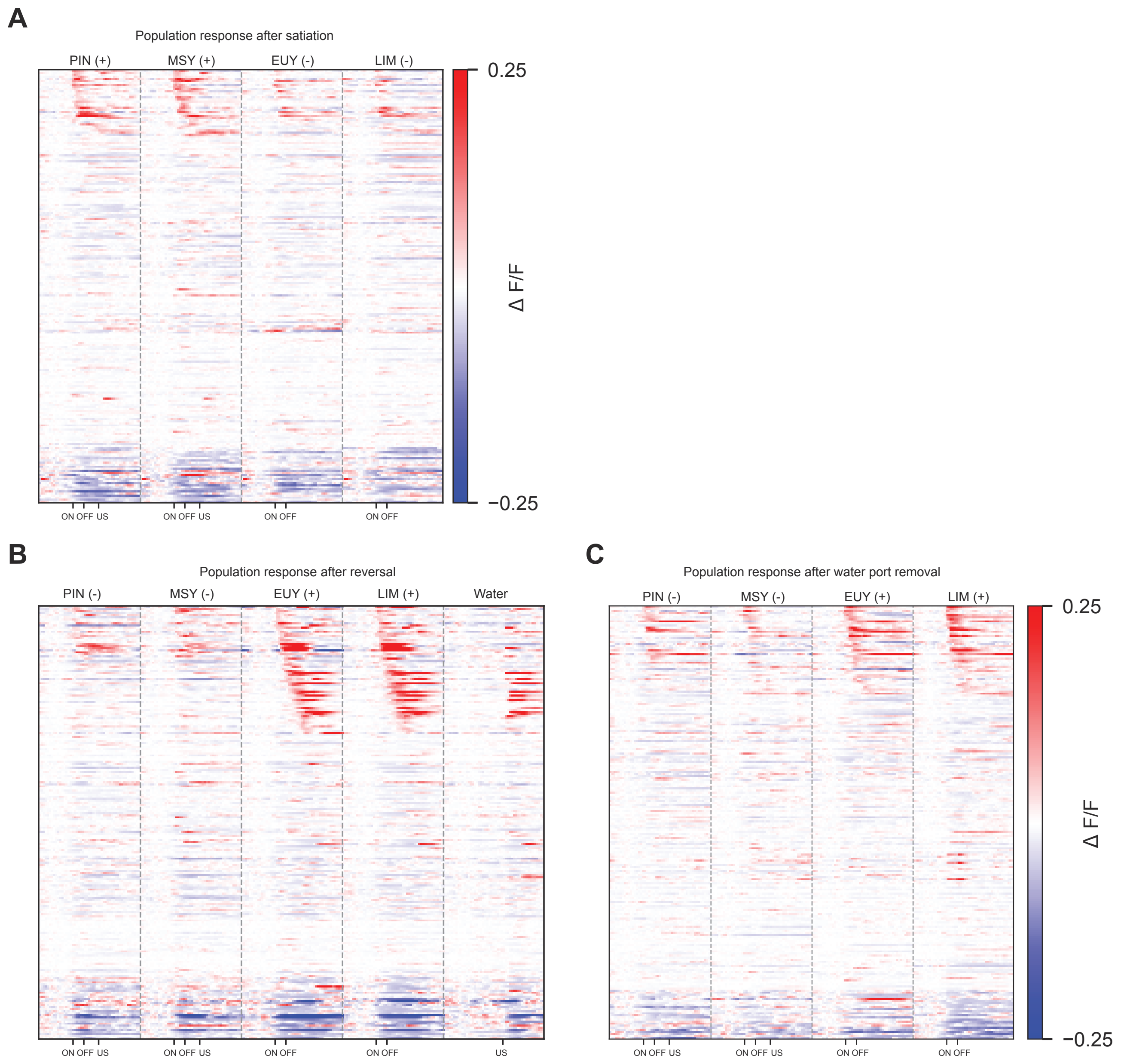
A
Population response after satiation
PIN (+)
MSY (+)
EUY (-)
LIM (-)
0.25
−0.25
Δ F/F
ON OFF US
ON OFF US
ON OFF
ON OFF
B
Population response after reversal
PIN (-)
MSY (-)
EUY (+)
LIM (+)
Water
ON OFF US
ON OFF US
ON OFF
ON OFF
US
C
Population response after water port removal
PIN (-)
MSY (-)
EUY (+)
LIM (+)
0.25
−0.25
Δ F/F
ON OFF US
ON OFF US
ON OFF
ON OFF

**Figure S3. All OFC imaging data during reversal learning, and during manipulations of internal state and external context. Related to Figures 2 and 3**

(A-C) PSTH of OFC responses pooled from all imaged mice (n=5) after satiation (A, corresponding to Figure 3E), after reversal learning (B, corresponding to Figure 3A), and after water port removal (C, corresponding to Figure 3G). Data were collected from the same mice as shown in Figure 2A and 2B. Satiation, reversal, and water port removal experiments were conducted in the order shown. As a consequence, during reversal and water port removal experiments, the rewarded odors were EUY and LIM.

### Figure S4

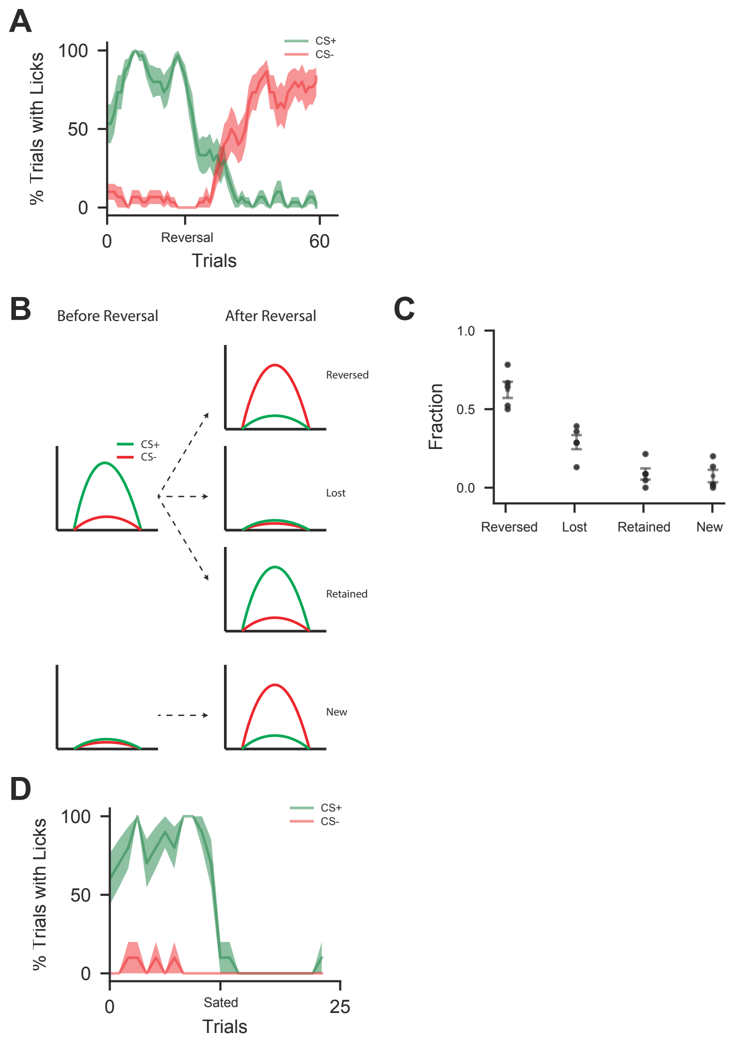
A
100
50
0
% Trials with Licks
CS+
CS-
0
Reversal
60
Trials
B
Before Reversal
After Reversal
CS+
CS-
Reversed
Lost
Retained
New
C
1.0
0.5
0.0
Fraction
Reversed
Lost
Retained
New
D
100
50
0
% Trials with Licks
CS+
CS-
0
Sated
25
Trials

**Figure S4. Quantification of behavioral and OFC imaging data during reversal learning and manipulations of internal state and external context. Related to Figure 3**

(A) Percentage of trials with anticipatory licking to CS+ (green) and CS- odors (red) during reversal learning, as in Figure 3A-D. Here and below, shading indicates ±1 SEM.

(B) Schematic of possible changes in CS+ responses in a neuron upon reversal learning. Responses may reverse, may be lost, or may remain unchanged during reversal learning. Alternatively, a neuron previously unresponsive to the old CS+ odors may acquire a new response to the new CS+ odors after reversal learning.

(C) Of all the neurons responsive to the CS+ odors during discrimination learning, 62% reversed their CS+ responses, 29% lost CS+ responses, and 9% retained CS+ responses during reversal learning. An additional 8% of neurons that were unresponsive to the odors rewarded during learning (old CS+) gained a response to the odors rewarded during reversal learning (new CS+). Error bars indicate mean ±1 SEM, and dots indicate individual animals.

(D) Percentage of trials with anticipatory licking to CS+ (green) and CS- odors (red) before and after satiation, as in Figure 3E-F.

Figure S5

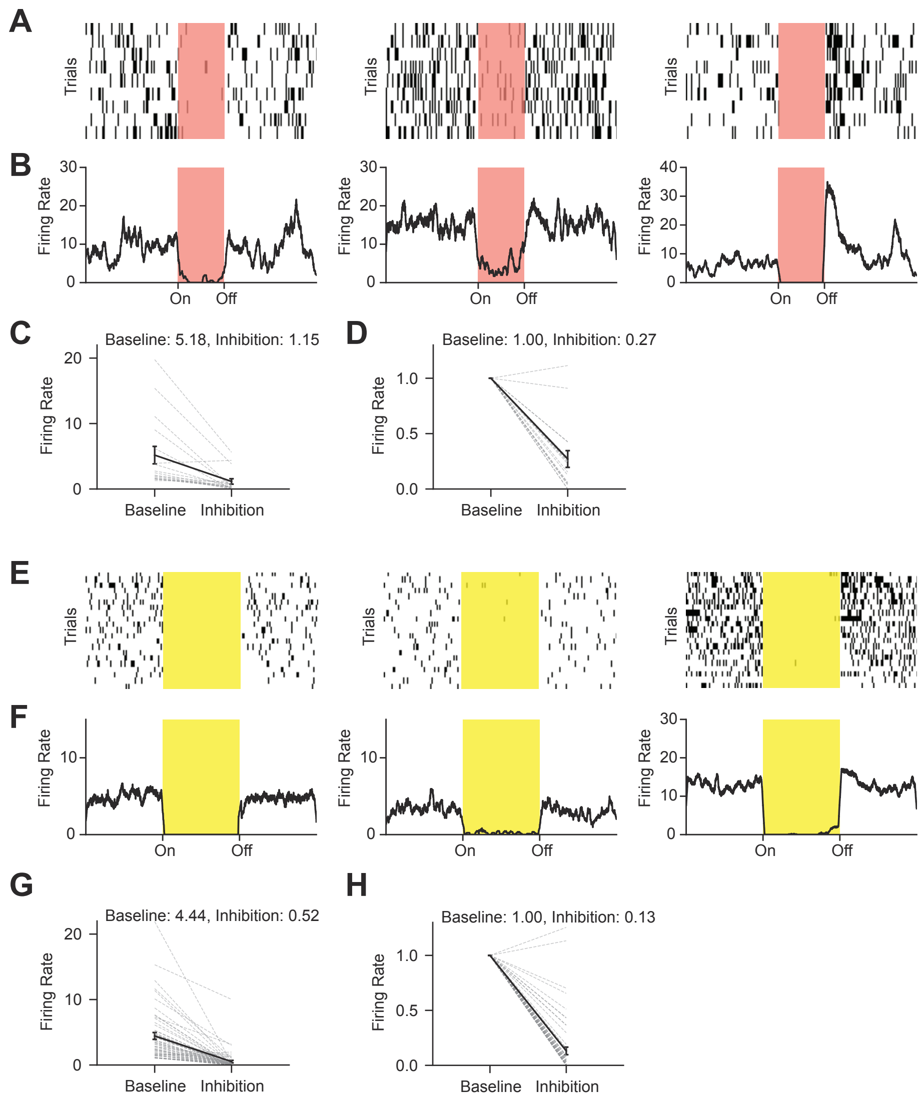
A
Trials
Trials
Trials
B
Firing Rate
Firing Rate
Firing Rate
On
Off
C
Baseline: 5.18, Inhibition: 1.15
Firing Rate
Baseline
Inhibition
D
Baseline: 1.00, Inhibition: 0.27
Firing Rate
Baseline
Inhibition
E
Trials
F
Firing Rate
On
Off
G
Baseline: 4.44, Inhibition: 0.52
Firing Rate
Baseline
Inhibition
H
Baseline: 1.00, Inhibition: 0.13
Firing Rate
Baseline
Inhibition

**Figure S5. Optrode recordings of OFC responses during OFC photostimulation in mice expressing Jaws or halorhodopsin. Related to Figures 4**

(A and B) Spike rasters (A) and firing rates (B) of 3 example OFC cells during photo-stimulation of OFC in mice expressing Jaws (n = 19 cells). Here and below, laser power was set to 5 mW out of fiber tip. See STAR Methods.

(C and D) Unnormalized (C) and normalized (D) firing rates of recorded neurons during baseline period and during OFC inhibition. Here and below, error bars indicate mean ±1 SEM.

(E and F) Spike rasters (E) and firing rates (F) of 3 example OFC cells during photo-stimulation of OFC in mice expressing halorhodopsin (n = 124 cells).

(G and H) Unnormalized (G) and normalized (H) firing rates of recorded neurons during baseline period and during OFC inhibition.

### Figure S6

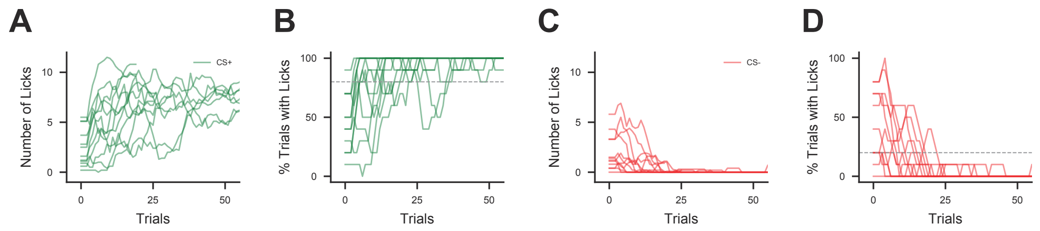
A
B
C
D
Number of Licks
% Trials with Licks
Trials
CS+
CS-
0
25
50
5
10
100

**Figure S6. Behavioral data in the discrimination phase of the two-phase learning task in head-fixed mice, related to Figures 5, 6, 7**

(A-D) Behavioral data acquired during head-fixed two-phase learning task for control mice without OFC or mPFC inhibition. Imaging and YFP cohorts are pooled (n=12 mice).

(A and B) Number of anticipatory licks (A) and percentage of trials with anticipatory licking (B) to CS+ odors during the discrimination phase of the two-phase task.

(C and D) Number of anticipatory licks (C) and percentage of trials with anticipatory licking (D) to CS- odors during the discrimination phase of the two-phase task.

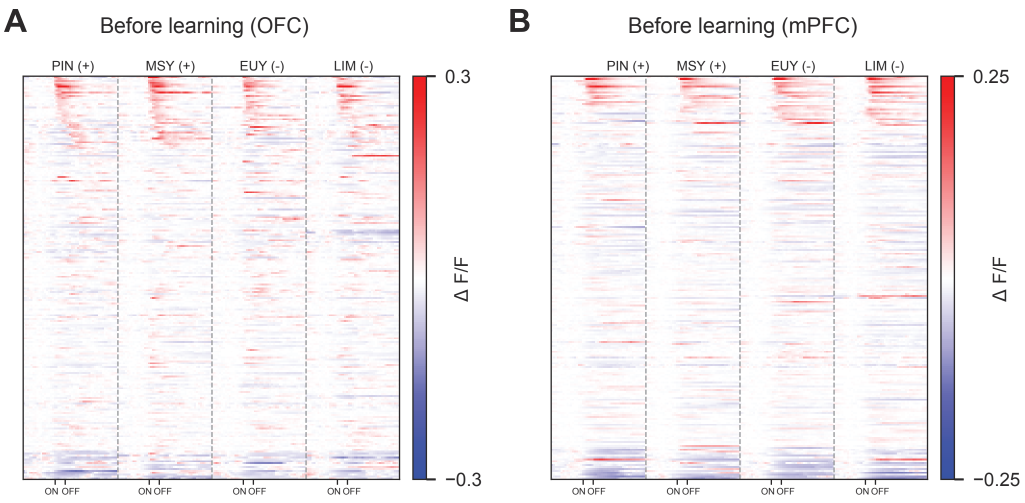
A
Before learning (OFC)
PIN (+)
MSY (+)
EUY (-)
LIM (-)
0.3
−0.3
Δ F/F
ON OFF
B
Before learning (mPFC)
0.25
−0.25

**Figure S7. OFC and mPFC population responses to odors before two-phase task learning in head-fixed mice. Related to Figure 6 and 7**

(A) PSTH of OFC responses to odors used in the discrimination phase of the two-phase task prior to any training. Responses are pooled across 4 mice, as for Figure 6.

(B) PSTH of mPFC responses to odors used in the discrimination phase of the two-phase task prior to training. Responses are pooled across 4 mice, as for Figure 7.

**TABLE S1, Related to STAR Methods**

| Brain Region | Lens and Virus Coordinates (mm) | Fiber Coordinates (mm) |
|---|---|---|
| Piriform | ML: 1.2<br>AP: 2.2<br>DV: -3.35 | NA |
| OFC | ML: 1.0<br>AP: 2.3-2.4<br>DV: -2.45 | ML: 1.0<br>AP: 2.3-2.4<br>DV: -2.1 |
| mPFC | ML: 0.4<br>AP: 1.65<br>DV: -2.05 | ML: 0.4<br>AP: 1.65<br>DV: -1.7 |

**TABLE S2, Related to STAR Methods**

| Odors | Abbr | Supplier | Identifier | Concentration (head-fixed) | Concentration (freely moving) |
|---|---|---|---|---|---|
| (1R)-(-)-Fenchone | fen | Sigma | 196436 | 2% | NA |
| (1R)-(+)-α-Pinene | pin | Sigma | 268070 | 2% | 1% |
| (R)-(+)-Limonene | lim | Sigma | 183164 | 2% | NA |
| 2-Phenylethanol | 2pe | Sigma | 77861 | 2% | NA |
| 3-Octanol | oct | Sigma | 218405 | 2% | NA |
| 4-Methylthiazole | 4mt | Sigma | 193925 | 2% | NA |
| Benzaldehyde | ben | Sigma | 418099 | 2% | NA |
| Eucalyptol | euy | Sigma | C80601 | 2% | NA |
| Geranyl acetate | ger | Sigma | 173495 | 2% | NA |
| Isopentyl acetate | iso | Sigma | 112674 | 2% | 1% |
| Methyl salicylate | msy | Sigma | M2047 | 2% | NA |
| Mineral Oil | oil | Sigma | M5904 | NA | NA |
| Ethyl Acetate | eth | Sigma | 270989 | NA | 1% |

**TABLE S3, Related to STAR Methods**

| Imaging Cohort | N | Brain Area | Experiment | Figures |
|---|---|---|---|---|
| A | 6 | Piriform | Single-phase discrimination learning, passive odor exposure [a] | 1 |
| B | 5 | OFC | Single-phase discrimination learning, reversal, internal state, external context [b] | 2, 3 |
| C | 3 | OFC | Discrimination (long-term) | 4G-I |
| D | 4 | OFC | Two-phase learning task | 6A-6L |
| E | 4 | mPFC | Two-phase learning task | 7A-7P |

a. 4 of 6 mice in cohort A were used for the passive odor exposure experiment

b. 4 of 5 mice in cohort B were used for the external context experiment (removal of water port).

**TABLE S4, Related to STAR Methods**

| Behavioral Cohort | N | Brain Area | Experiment | Virus | Source | Concentration (vg/mL) | Volume (nL) |
|---|---|---|---|---|---|---|---|
| F | 4 | OFC | Discrimination, single-phase, head-fixed | AAV-CamKII-Jaws-KGC-GFP-ER2 | UPenn Vector Core | 5 x 10^12 | 300 |
| G | 5 | OFC | Discrimination, single-phase, head-fixed | AAV-hSyn-eNPHR3.0-EYFP | UNC Vector Core | 5 x 10^12 | 1000 |
| H | 4 | OFC | Discrimination, single-phase, head-fixed | AAV-hSyn-EYFP | UNC Vector Core | 3.3 x 10^12 | 300 |
| I | 6 | OFC | Pretraining, two-phase, head-fixed | AAV-CamKII-Jaws-KGC-GFP-ER2 | UPenn Vector Core | 5 x 10^12 | 300 |
| J [a] | 4 | OFC | Pretraining and discrimination, two-phase, head-fixed | AAV-hSyn-EYFP | UNC Vector Core | 3.3 x 10^12 | 300 |
| K | 4 | OFC | Discrimination, two-phase, head-fixed | AAV-CamKII-Jaws-KGC-GFP-ER2 | UPenn Vector Core | 5 x 10^12 | 300 |
| L | 5 | OFC | Pretraining, two-phase, freely moving | AAV-hSyn-eNPHR3.0-EYFP | UNC Vector Core | 5 x 10^12 | 1000 |
| M | 6 | OFC | Pretraining, two-phase, freely moving | AAV-hSyn-EYFP | UNC Vector Core | 3.3 x 10^12 | 1000 |
| N | 5 | OFC | Discrimination, two-phase, freely moving | AAV-hSyn-eNPHR3.0-EYFP | UNC Vector Core | 5 x 10^12 | 1000 |
| O | 4 | OFC | Discrimination, two-phase, freely moving | AAV-hSyn-EYFP | UNC Vector Core | 3.3 x 10^12 | 1000 |
| P | 5 | mPFC | Pretraining, two-phase, freely moving | AAV-hSyn-eNPHR3.0-EYFP | UNC Vector Core | 5 x 10^12 | 1000 |
| Q | 7 | mPFC | Pretraining, two-phase, freely moving | AAV-hSyn-EYFP | UNC Vector Core | 3.3 x 10^12 | 1000 |
| R | 4 | mPFC | Discrimination, two-phase, freely moving | AAV-hSyn-eNPHR3.0-EYFP | UNC Vector Core | 5 x 10^12 | 1000 |
| S | 4 | mPFC | Discrimination, two-phase, freely moving | AAV-hSyn-EYFP | UNC Vector Core | 3.3 x 10^12 | 1000 |

a. cohort J were used as controls for both pretraining and discrimination learning in the two-phase task in head-fixed animals.

**MOVIE S1. Recordings of Piriform, OFC, and mPFC**

Example video showing calcium activity recorded in piriform, OFC, and mPFC in three separate mice after motion correction (STAR Methods). Images were scanned at approximately 4.5 Hz, and playback framework is 30 frames per second (7x speed up).

**MOVIE S2. Tracking of OFC neurons across 9 imaging days**

Example video showing all tracked ROIs overlaid on the mean intensity images for 9 days of imaging in OFC. The shapes of the ROIs are updated to accommodate differences in cell shapes on each imaging day (STAR Methods). Same field of view as shown in the OFC recording in Video 1.
